# A multiplex microchamber diffusion assay for the antibody-based detection of microRNAs on randomly ordered microbeads

**DOI:** 10.1101/2021.03.19.436219

**Authors:** Christiane Geithe, Bo Zeng, Carsten Schmidt, Franziska Dinter, Dirk Roggenbuck, Werner Lehmann, Gregory Dame, Peter Schierack, Katja Hanack, Stefan Rödiger

## Abstract

**Background:** MicroRNAs (miRNAs) are small, conserved, noncoding RNAs regulating gene expression that functions in RNA silencing and post-transcriptional regulation of gene expression. Altered miRNA profiles have been implicated in many human diseases, and due to their circulating abilities, they have excited great interest in their use as clinical biomarkers. The development of innovative methods for miRNA detection has become of high scientific and clinical interest.

**Methods:** We developed a diffusion-driven microbead assay and combined it with an antibody-based miRNA detection. The diffusion process was carried out in two different approaches a) co-diffusion of miRNA and antibodies (termed diffusion approach I, DAI) and b) diffusion of miRNA in an antibody-saturated environment (DAII). In both approaches, neutravidin-coated microbeads were loaded with specific biotinylated DNA capture probes, which targets either miR-21-5p, miR-30a-3p or miR-93-5p. The miRNAs were time- and dose-dependently detected in a diffusion microchamber by primary anti-DNA:RNA hybrid and fluorescence-labeled secondary antibodies using our in-house developed inverse fluorescence microscope imaging platform VideoScan.

**Results:** Our assay offers the advantage that several target molecules can be detected simultaneously and in real-time in one reaction environment (multiplex), without any amplification steps. We recorded the diffusion process over a period of 24 h and found that the reaction was almost completed after 2 h. The specificity of the assay was 96.7 % for DAI and 92.3 % for DAII. The detection limits were in a concentration range of 0.03-0.43 nM for DAI and 0.14-1.09 nM for DAII, depending on the miRNA.

**Conclusion:** The miRNAs are successively exposed to the capture probe-loaded randomly ordered microbeads (p value of CSR 0.23-0.96), which leads to microbeads that become saturated with the target molecules first in front rows. Non-bonded miRNAs continue to diffuse further and can therefore subsequently bind to the microbeads with free binding sites. Our detection principle differs from other microbead assays, in which all microbeads are simultaneously mixed with the sample solution, so that all target molecules bind equally distributed to the microbeads, resulting in an averaged signal intensity.

## Introduction

MicroRNAs (miRNAs) are short (20-22 nucleotides), highly conserved, noncoding RNAs, which regulate gene expression post-transcriptionally by targeting mRNAs, thus leading to degradation of mRNA or inhibition of translation. They have been identified as biomolecules that are involved in the development of many human diseases, such as cancer, inflammatory diseases, neurodegenerative diseases or immune-related diseases [1, 2].

The miRNAs were also associated with heart disease [3–5]. Heart disease is one of the most widespread causes of death in western countries. The dilated cardiomyopathy (DCM) is a major contributor to heart disease. DCM has a mortality rate of 50 % within five years and is the most common cause of heart transplantation. Characteristics for DCM are the strong volume increase (dilatation) of the left and right ventricles and a homogeneous contractile dysfunction of the myocardium with the consequence of a restricted systolic function [6–9].

Due to the ability of miRNAs to circulate in body fluids such as blood, plasma or serum, and their stability therein, they emerged as potent biomarkers for diagnosis and prognosis [10, 11]. Altered miRNA profiles associated with certain diseases can be identified by technologies such as high-throughput sequencing, microarrays or quantitative PCR (qPCR) analysis [11–13]. It is also generally accepted that amplification-based systems can be biased, as amplification can lead to a shift in certain target molecules [14]. To reduce technical demands, costs and hands-on time, the development of alternative methods for miRNA detection has become a field of intensive research and clinical interest. Amplification-free detection systems in particular are repeatedly the subject of research work. In the assay described by Kappel *et al.* [15], for example, biotinylated complementary capture probes are hybridized directly with miRNAs and then the resulting hybrids are bound to a solid phase. Optical detection is performed with an anti-DNA:RNA hybrid antibody labeled with an acridinium ester, resulting in subsequent enzymatic signal amplification [15]. In contrast, the nanostring technology uses two complementary probes for hybridization, a target specific biotinylated capture probe and a target specific barcoded fluorescence reporter probe. The unique target-probe complex is immobilized and aligned on the imaging surface for the fluorescence readout [16].

Optical methods are common and widely used for the detection of biomolecules or especially miRNAs. We used our in-house developed *VideoScan* platform [17] as a readout platform, which is based on a fully automated multispectral fluorescence microscope. For this, we have developed multiplex assays for the imaging platform to detect and quantify biomolecules including short protein or DNA sequences [18] and (auto)antibodies [19]. The platform is also suitable for real-time measurements such as qPCRs, isothermal amplification and melting curve analysis [20, 21]. Microbead-based assays offer the advantage that several target molecules can be detected and quantified simultaneously in a reaction environment (multiplex). Planar microarrays have disadvantages regarding the mechanisms of surface interactions. In particular, the mass transport on the surface is limited [22]. Especially for the detection of short single-stranded molecules there are a number of detection principles based on molecular beacons and FRET probes [23]. Their detection principle is based on microbeads that have specific capture probes on their surface to which the target molecules can bind. All microbeads are simultaneously (bulk) mixed with the sample solution, so that in principle all target molecules can bind equally distributed to the microbeads. The binding event is then quantified by determining the surface signal intensity of all microbeads.

An alternative approach is that the target molecules are successively exposed to the capture probe-loaded microbeads. One can postulate as a hypothesis that in this approach the capture probe-loaded microbeads are saturated with the target molecules. Non-bonded target molecules can therefore subsequently bind to the microbeads with free binding sites. Based on this hypothesis, we have developed a microchamber in which the target molecules (miRNAs) diffuse through a field of capture probe-loaded microbeads and govern with their binding partners. The aim of this study was to design an antibody-based amplification-free assay that does not require labelling of miRNAs and can detect multiple molecules simultaneously (multiplex) at a defined temperature (isothermal). A further criterion was a stepwise and real-time resolved visualization of the binding process between the target nucleic acids and their capture molecules.

## Methods

### Assembly of the diffusion microchamber

We used hybriwell chambers (16 wells, 7 mm x 7 mm x 0.05 mm; RD477991, Grace Bio-Labs, Oregon, USA) that were customized according to our needs. To make the hybriwell chamber suitable for long-time experiments, we have modified them with reaction reservoirs at the inlets and outlets (Fig. 1). First, the adhesive hybriwell chamber (Fig. 1A) was glued to a glass slide. To modify the chamber, PCR reaction vessels were cut to a size of about 6 mm by cutting off the upper and lower ends using a custom-made cutting device (Fig. 1B, C). The resulting truncated hollow cones were then attached around the chamber inlets and outlets using a mixture of UV adhesive and carbon black. The carbon black was used to prevent the auto-fluorescence of the UV adhesive. The adhesive mixture was cured by UV irradiation (2 x 3 min at 312 nm) in a UV cross-linker (BIO-Link BLX-E, Vilber Lourmat, Germany). The modified microchamber consists of 16 cavities, each with a reservoir at the inlet and outlet (Fig. 1D). The attached reservoirs were used to seal the reaction chamber airtight to prevent evaporation or formation of air bubbles that could impair the diffusion process.

**Figure 1.**
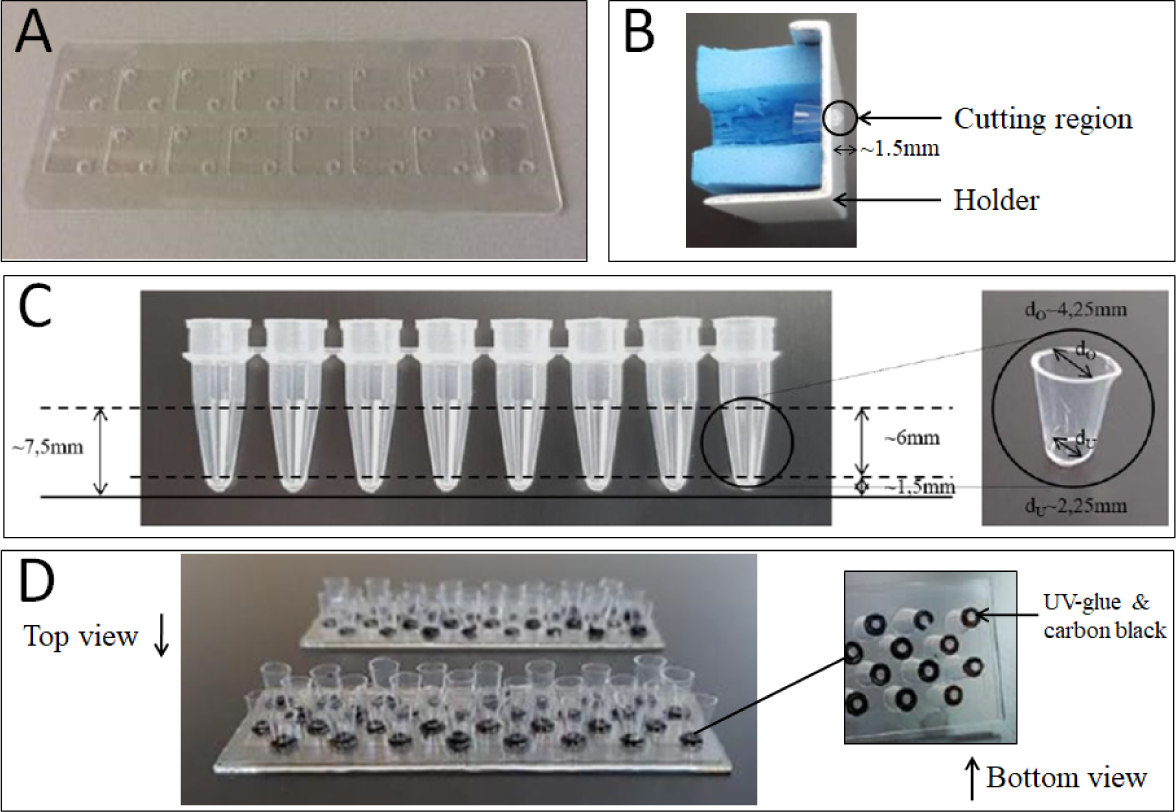
Modification of the hybriwell chamber. (A) GraceBio-Labs hybriwell chamber, (B) custom-made cutting device for precisely cutting the reaction vessels, (C) cutting of the reaction vessels to size, (D) modified microchamber from top and bottom.

### Immobilizing of miRNA-specific capture probes on microbeads

#### Microbeads

We used three different carboxylated poly(methyl methacrylate) (PMMA) microbead (MB) populations (PolyAn GmbH, Germany) which differ in size (MB1: 11 µm, MB2: 11.5 µm, MB3: 12 µm) and a specific ratio of two fluorescence dyes (rhodamine 6G, excitation wavelength Ex 520 nm, emission wavelength Em 560 nm; coumarin 6, Ex 460 nm, Em 500 nm).

### Covalent coupling of neutravidin

The size/dye-encoded microbeads were coated with neutravidin (Thermo Fisher Scientific, Germany) by covalent coupling using 1-ethyl-3-(3-dimethylaminopropyl) carbodiimide hydrochloride (EDC; Karl Roth, Germany) cross-linker according to [24]. Neutravidin was used because it has a high temperature stability and a high binding constant [25]. In brief, 1.5×10^5^ microbeads were first washed with 200 µL 0.1 M 2-(N-morpholino) ethanesulfonic acid (MES, pH 4.5; Karl Roth, Germany) and incubated for 30 min at 28°C and 1,200 rpm with 100 µL EDC solution (25 mg/mL EDC dissolved in MES). The microbeads were then washed with 200 µL 0.05 x phosphate buffered saline (PBS) and incubated for 3 h at 28 °C and 1,200 rpm with 50 µL neutravidin solution (60 µg/mL dissolved in 0.05 x PBS). This was followed by three washing steps with Tween-20 containing Tris-buffered saline (TBS-T; 50 mM Tris, 154 mM NaCl, 0.01 % Tween-20, pH 7.4). Coupling of miRNA-specific capture probes Dual-biotinylated miRNA-specific DNA capture probes (50 nM; complementary DNA sequence to the respective target) for hsa-miR-21-5p (5’-TCA ACA TCA GTC TGA TAA GCT A-dual-Biotin-3’), hsa-miR-30a-3p (5’-GCT GCA ACA TCC GAC TGA AAG-dual-Biotin-3’) or hsa-miR-93-5p (5’-CTA CCT GCA CGA ACA GCA CTT TG-dual-Biotin-3’; Biomers.net, Germany) were each immobilized on a different neutravidin-coated microbead population. The reaction was conducted in a volume of 50 µL for 15 min at room temperature under vigorous shaking (1,200 rpm) in TBS-T followed by three times wash with PBS containing Tween-20 (PBS-T; 137 mM NaCl, 2.7 mM KCl, 10 mM Na_2_HPO_4_, 2 mM KH_2_PO_4_, 0.01 % Tween-20, pH 7.4). Afterwards, all three capture probe-loaded microbead populations were bulk mixed (3-plex) in comparable proportions (approx. 1,000 beads per population).

### Sample preparation for the multiplex microbead-based miRNA detection by antibodies in a diffusion microchamber

#### Diffusion approach I – co-diffusion of miRNAs and antibodies

The procedure of sample preparation is illustrated in Figure 2. First, a 7 µL suspension containing three different microbead populations (3-plex, approx. 1,000 beads per population) with specific DNA capture probes (cp) for either hsa-miR-21-5p, hsa-miR-30a-3p or hsa-miR-93-5p were added from one side of the microchamber and distributed randomly over the entire bottom surface (Fig. 2A). The miRNAs (0.05 nM, 0.1 nM, 0.5 nM, 1 nM, 5 nM, 10 nM or 50 nM hsa-miR-21-5p: 5’-UAG CUU AUC AGA CUG AUG UUG A-3’, hsa-miR-30a-3p: 5’-CUU UCA GUC GGA UGU UUG CAG C-3’ or hsa-miR-93-5p: 5’-CAA AGU GCU GUU CGU GCA GGU AG-3’; Biomers.net, Germany) to be detected, the primary antibody (2.0µg/mL; anti-DNA:RNA hybrid, clone S9.6; monoclonal, mouse, IgG2ak, Merck Millipore, Germany) and the secondary fluorescence-labeled anti-mouse IgG antibody (2.5 µg/mL; goat anti-mouse IgG, Atto647N, Sigma Aldrich, Germany) were mixed in one batch and offered in a volume of 2 µL to the beads from the opposite side. Both sides of the chamber were then sealed with 10 µL mineral oil using the reservoirs. All components (capture probe-loaded microbeads, miRNAs and antibodies) were diluted with PBS-T buffer. The entire diffusion process was carried out in PBS-T.

**Figure 2.**
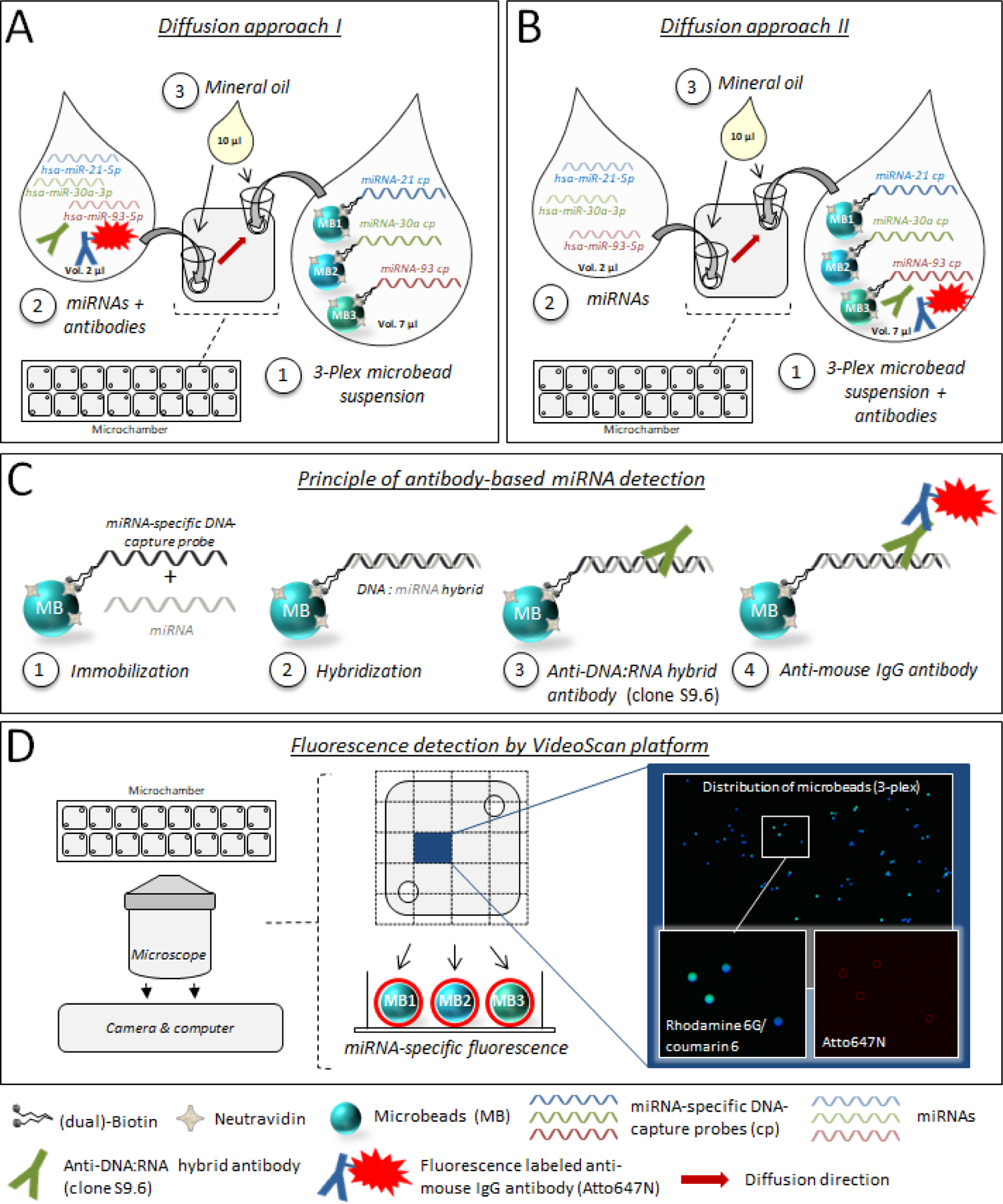
Multiplex microchamber diffusion assay workflow. Shown are two different experimental approaches: (A) “co-diffusion of miRNAs and antibodies” and (B) “miRNA diffusion in an antibody-saturated environment”. In the first approach, a bulk mixed suspension of three microbead populations, each carrying a different specific DNA capture probe for miRNAs, was added from one side of the diffusion microchamber and the miRNAs (either as a mixture or individually) were applied together with the antibodies from the opposite side. In the second one, only the miRNAs (either as a mixture or individually) were added into the microchamber in which the microbeads and antibodies were already distributed. (C) Principle of miRNA detection with an anti-DNA:RNA hybrid antibody. (D) miRNA-specific fluorescence readout with the VideoScan platform. Shown are VideoScan false colour images of the distributed size/dye-encoded microbeads (3-plex) as well as miRNA-specific fluorescence signals (Atto647N) which become visible as a red corona around the beads.

### Diffusion approach II – miRNA diffusion in an antibody-saturated environment

In contrast to the first approach, the capture probe-loaded microbeads (3-plex; see above) were added together with the anti-DNA:RNA hybrid antibody (2.0 µg/mL), and the secondary antibody (2.5 µg/mL) from one side of the microchamber (Fig. 2B). The respective miRNAs were offered to the microbead/antibody environment from the opposite site, and both sides were subsequently sealed with mineral oil.

### Principle of antibody-based miRNA detection

MiRNA-specific DNA capture probes were firstly immobilized on neutravidin-coated microbeads. The respective miRNA hybridizes with its capture probe, leading to the formation of a DNA:miRNA hybrid, which was then recognized by an anti-DNA:RNA hybrid antibody and a fluorescent secondary antibody (Fig. 2C).

The specificity of the anti-DNA:RNA hybrid antibody towards DNA:miRNA hybrids was shown in several studies. Complementary and thus perfect matched miRNAs to the capture probe lead to a perfect antibody binding (Fig. 3A), whereas single-base, multiple-base miRNA mismatches or non-complementary miRNA lead to impaired or no antibody binding (Fig. 3B) [15, 26].

**Figure 3.**
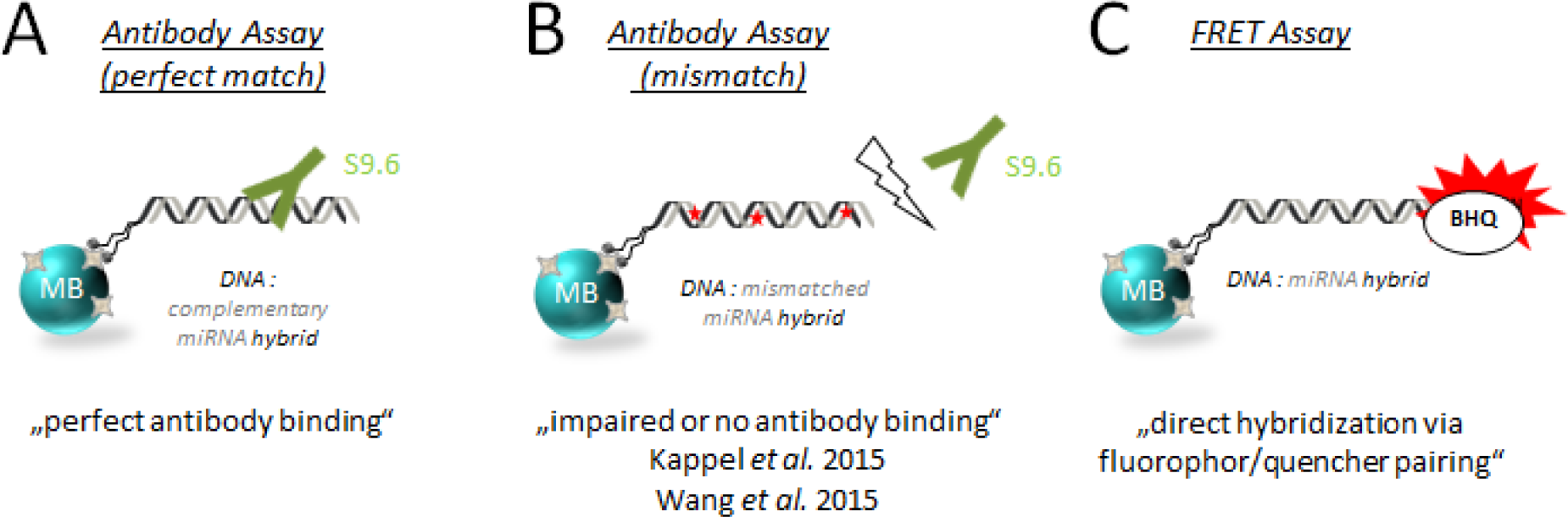
MiRNA detection methods. (A) Recognition of perfectly matched DNA:miRNA hybrids by the anti-DNA:RNA hybrid antibody clone S9.6. (B) Single-base or multiple-base mismatches between the miRNA-specific DNA capture probe and the miRNA lead to impaired or no antibody binding. The red stars indicate mismatches. (C) FRET-based direct hybridization of a fluorophore-labeled miRNA-specific DNA capture probe with a quencher-labeled miRNA. This FRET system was used for melting temperature determination.

### Fluorescence detection by *VideoScan* platform

The fluorescence signals were monitored at room temperature (24 °C ± 2 °C) and different time points (5 min, 40 min, 100 min, 160 min, 220 min, 280 min, 340 min and 24 h) by the *VideoScan* platform, which automatically records 20 individual images per microchamber well. The platform is able to distinguish between different microbead populations due to their size and specific dye ratios of rhodamine 6G and coumarin 6 [17]. Microbeads of the same population have the same size, the same fluorescence dye ratio and a certain degree of carboxylation. Microbead-associated, and thus miRNA-specific, fluorescence signals (Atto647N) become visible as a red corona around the beads (Fig. 2D).

### Characterization of DNA:miRNA hybrids

The optimal temperature is important to minimize the contribution of mismatch nucleic acid species [27]. The melting temperatures of the DNA:miRNA hybrids on the surfaces of the microbeads were determined with the *VideoScan* heating and cooling unit (HCU) [17] and then evaluated with the MBmca package (v. 0.0.3-5) [20] using the R GUI /IDE RKWard version 0.7.1z+0.7.2+devel1 [28]. For this purpose, the DNA:miRNA hybrids were first formed on the microbeads by using synthetic DNA-fluorophor/miRNA-quencher pairs (Fig. 3C) (miRNA-21: 5’-Atto647N-TCA ACA TCA GTC TGA TAA GCT A-dual-Biotin-3’/ 5’-UAG CUU AUC AGA CUG AUG UUG A-BHQ2-3’; miRNA-30a: 5’-Atto647N-GCT GCA ACA TCC GAC TGA AAG-dual-Biotin-3’/ 5’-CUU UCA GUC GGA UGU UUG CAG C-BHQ2-3’; miRNA-93: 5’-Atto647N-CTA CCT GCA CGA ACA GCA CTT TG-dual-Biotin-3’/ 5’-CAA AGU GCU GUU CGU GCA GGU AG-BHQ2-3’, Biomers.net, Germany). For the determination of the melting temperature, the temperature was then continuously increased from 30 °C to 80 °C (1°C/step) and the change in fluorescence intensity was recorded over time in PBS-T buffer. The melting temperatures were determined from the maximum of the first negative derivative (Table 1). Calculated melting temperatures and some other thermodynamic properties for the miRNAs used in this study are given in Table S1.

**Table 1:** Determined melting temperatures of DNA:miRNA hybrids in the microbead-based FRET assay.

| **miRNA** | **melting temperature**<br>**(experimental)**<br>**$T_M$ [°C]** |
| --- | --- |
| miR-21-5p | 54.8 |
| miR-30a-3p | 56.5 |
| miR-93-5p | 64.3 |

### Data analysis

#### Data preprocessing, normalization and visualization

The data were analyzed using analysis pipelines implemented in *Python* (3.+) to specifically analyze point pattern datasets. The raw *VideoScan* data (comma separated values) were reformatted to be read by the *Python* package *pandas* (v. 0.23.0). After extracting the fluorescence signal of population-specified microbeads, a threshold value using 99 % quantile rule was calculated to detect microbeads with excessively high intensities at each time point. Those disproportionately fluorescent microbeads were assigned to the threshold value. Each microbead-specific fluorescence of each time point (5 min - 24 h) was normalized to the maximum fluorescence of the highest miRNA concentration (50 nM) of the 24 h measurement. The normalized signals were used to create the contour plots for distinct time points as well as miRNA concentrations using the *Python* package *matplotlib* (v. 2.2.2). To observe the number of positive microbeads at each time point, we used the 68–95–99.7 rule. Fluorescent signal intensities higher than mean + 3σ of the PBS-T control over time were considered as positive. From that the means, given as percent, were presented as bar plots. The fluorescence intensities of those positive microbeads were plotted using boxplots.

#### Dose-response curve fitting

For the diffusion kinetics, the change over time (dose-response curve) of positive microbead percentages and the fluorescence intensities of those positive microbeads were fitted for each miRNA concentration by a four-parameter log-logistic function (Equation 1) [29],

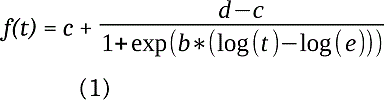

where *t* is the time, *f(t)* is the fluorescence intensity, *b* is the hill slope, *c* is the minimum intensity, *d* is the maximum intensity, *e* is the half maximal effective coefficient (*EC*_50_).

When making the dose-response curve fitting, we converted the x-axis dose values to logarithmic scale (log10(1e2 * dose). This will be helpful to plot the fitted curve into sigmoid shape, and this will make the relation between dose and response to be linear and therefore we can better choose the optimized dose values such as *EC_50_*. The logarithm conversion values for time and concentration are shown in (Table S2).

**Table 2.**
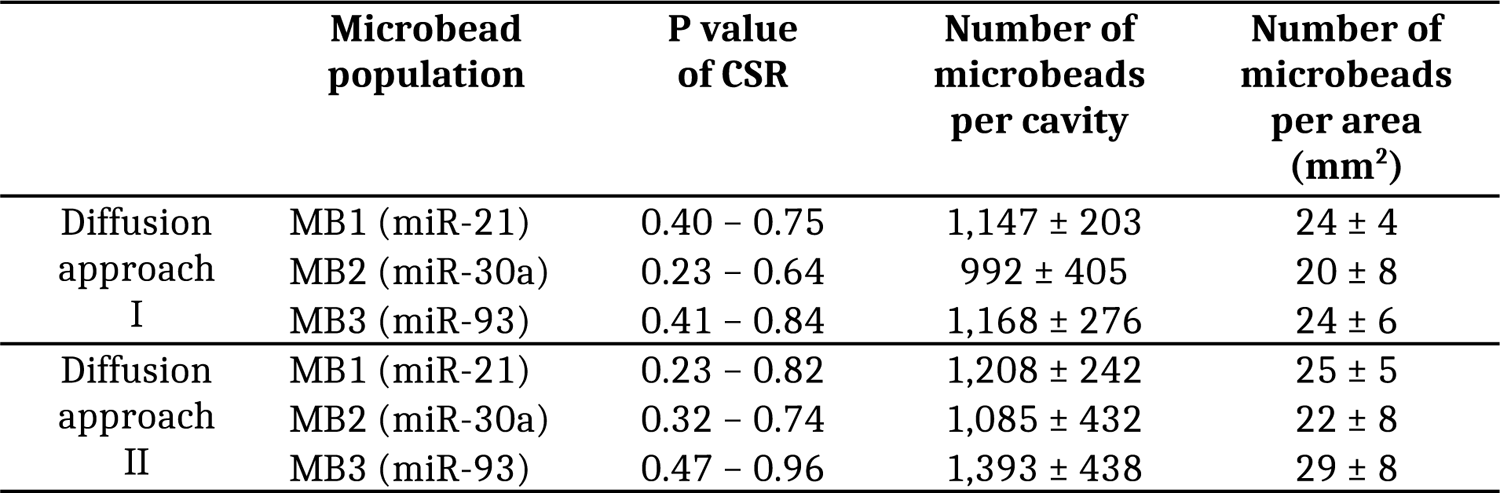
Test for random distribution of microbeads in the microchamber (p value of CSR), number of microbeads per area (mm^2^) and total number of microbeads per cavity of the microchamber.

For each of the two diffusion approaches, 3 biological replication experiments were conducted to verify the sensitivity of the detection system. Therefore, for each diffusion approach the mean (n=3) and standard deviation was calculated, and the values were used to make the curve fitting for fluorescence intensity and the number of positive microbeads.

#### Quantification point estimation by Second Derivative Maximum method

In mathematics, the first derivative can tell us the function *f(x)* is increasing (positive) or decreasing (negative), and the second derivative tells us the change rate, so the second derivative maximum (*SDM*) value could be interpreted as the inflection point with a maximum slope [30]. In this paper, with the use of the *diff* function from the *Python sympy* package, we calculated the *SDM* values for the time-and concentration-dependent increase curves of the number of positive microbeads and of the fluorescence intensities. We want to observe where the sigmoid curve gives the maximum turning point, and we can also compare the maximum turning points among different concentrations or time conditions.

#### Complete spatial randomness (CSR)

To test the randomness of the point pattern (spatial distribution) for each of the three microbead populations in a cavity of the microchamber, all cavities were judged against the hypothesis of complete spatial randomness (CSR). Therefore, we applied quadrat statistics to test individually the point pattern for all the microbead populations’ distribution in each cavity of the microchamber of several assays (mean *±* standard deviation for diffusion approach I, n=6 and diffusion approach II, n=6). For *P* < 0.05, we infer that the spatial distribution of our microbeads come not from a CSR process. This means that microbeads are not randomly distributed in the microchamber cavity. In addition to this we calculated the number of microbeads per area (unit) and the total number of microbeads per cavity of the chamber.

To investigate dependence between the points in a point pattern for each of the three microbead populations in the cavity, we estimated the cumulative distribution function *G* [31, 32] of the nearest-neighbor distance for the typical point pattern X (Equation 2):

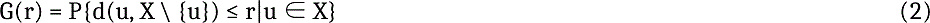

where *r* is the distance argument, *u* is an arbitrary location, and *d*(u, X \ {u}) is the shortest distance from *u* to the point pattern *X* excluding *u* itself. For inferential purposes, the estimate of *G* (empirical) is usually compared to the true value of *G* (theoretical) for a completely random (Poisson) point process, if empirical *G(r)* is bigger than theoretical *G(r)*. Thus the nearest-neighbour distances in the point pattern are shorter than for a Poisson process. This suggests a clustered pattern (where points tend to be close together), on the contrary it suggests a regular pattern (where points tend to avoid each other). For each of the three populations in the study, we calculated the experimental and theoretical *G(r)* and found these two plots are very close, which suggests our microbeads are independently distributed in a poisson process pattern.

## Results and Discussion

### Diffusion microchamber

We developed a microchamber diffusion assay, which enables the multiplex detection of miRNAs on microbeads by antibodies in a small reaction environment. We used a setup, where the microbeads are randomly ordered (*P* value of CSR > 0.05) (Table 2). Between 1,000 to 1,400 microbeads per population, which refer to 20-30 microbeads per mm^2^, were used in a single microchamber (Table 2). This number led to an optimal and random distribution of the microbeads in the diffusion microchamber.

In this assay, the miRNAs to be detected diffuse through a field of DNA capture probe-loaded microbeads, hybridize with their respective binding partners and form a DNA:miRNA hybrid. The hybrid is then detected by binding a primary antibody that specifically recognizes DNA:RNA hybrids, followed by binding a secondary fluorescence-labeled anti-mouse antibody that enables the detection of fluorescence signals by the *VideoScan* platform. A microbead-associated fluorescence signal can only be detected if the respective miRNA, the primary anti-DNA:RNA hybrid antibody and also the secondary, fluorescence-labeled antibody have found their particular target molecules.

The diffusion assay was performed in two different approaches. In the first approach, the miRNAs diffuse together with the primary and secondary antibodies through a field of microbeads loaded with capture probes (diffusion approach I - co-diffusion of miRNA and antibodies). In the second approach, however, only the miRNAs diffuse. Here, the two antibodies are already present in the microchamber along with the microbeads carrying the capture probes (diffusion approach II - diffusion of miRNAs in an antibody-saturated environment). We tested two approaches, in order to not lose any information e.g. regarding sensitivity or specificity.

### Specificity of the microchamber diffusion assay

To determine the specificity of our assay, we have carried out several experiments. The assay was first performed according to the principle of diffusion approach I (co-diffusion of miRNA and antibodies) with three selected miRNAs. In order to find out whether the complementary capture probes specifically bind their corresponding miRNAs, a 3-plex microbead mixture consisting of the populations MB1 with capture probe for miR-21 (MB1:miR-21_CP), MB2 with capture probe for miR-30a (MB2:miR-30a_CP) and MB3 with capture probe for miR-93 (MB3:miR-93a_CP) was inserted into the diffusion microchamber. The respective miRNA to be detected was added together with the anti-DNA:RNA hybrid/secondary antibody mixture and the diffusion was recorded over time (Fig. 4). The diffusion of the three single miRNAs hsa-miR-21-5p (Fig. 4A), hsa-miR-30a-3p (Fig. 4B) or hsa-miR-93-5p (Fig. 4C) showed only fluorescence signals on the microbeads carrying the respective complementary capture probe. This indicated no unspecific binding of miRNAs (assay specificity of 96.7 %). Control experiments, in which only the primary and secondary antibody without any miRNA (PBS-T control) was added to the capture probe-loaded microbeads (3-plex) showed no unspecific signals.

**Figure 4.**
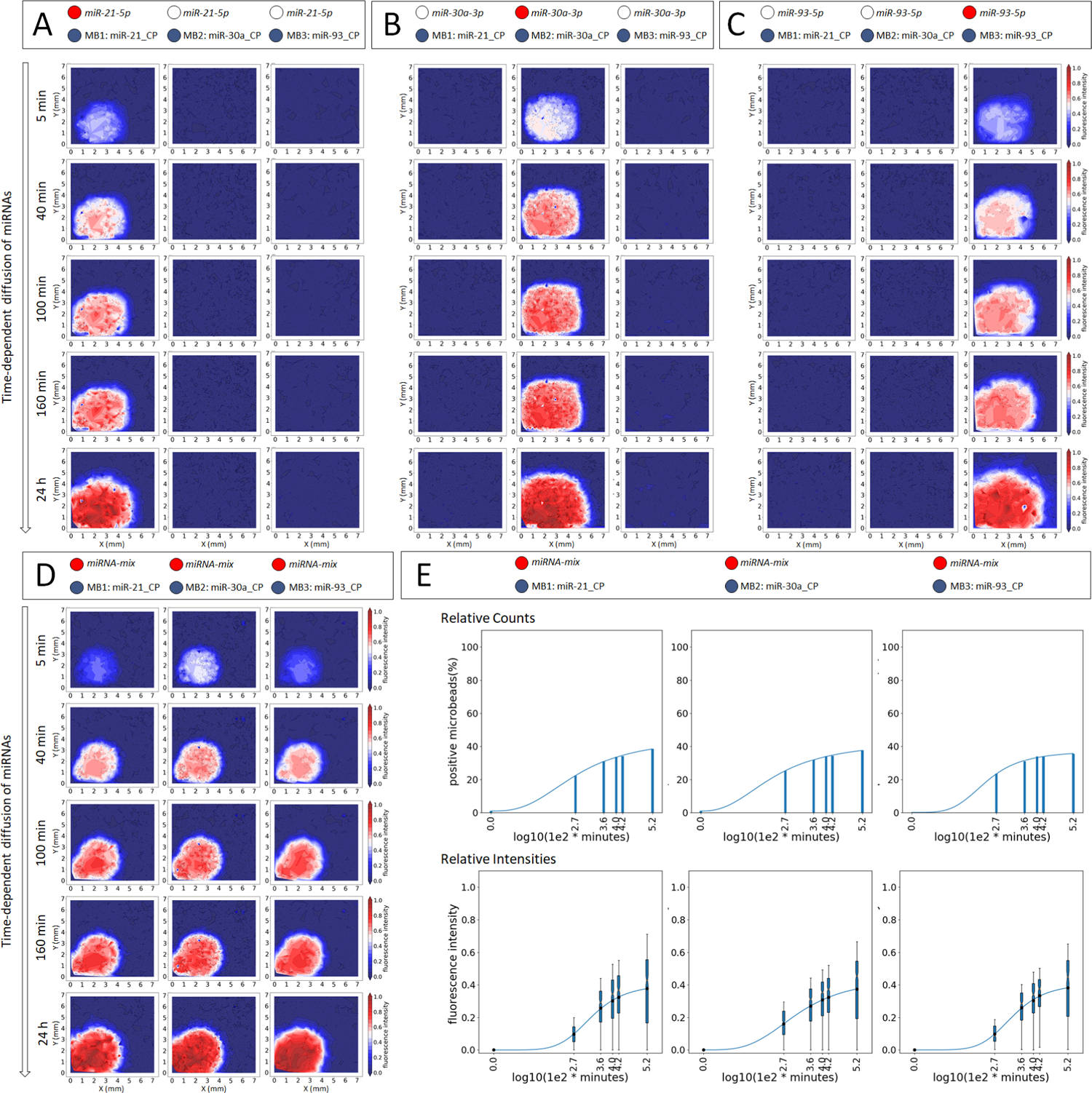
Multiplex diffusion-driven antibody-based detection of miRNAs in a microchamber. Time-dependent co-diffusion of (A) hsa-miR-21-5p (50 nM), (B) hsa-miR-30a-3p (50 nM), (C) hsa-miR-93-5p (50 nM) and antibodies through a field of miRNA-specific capture probe-loaded microbeads (3-plex). The signal intensities of each microbead population are individually displayed as contour plots at different time points. (D) Co-diffusion of the miRNA mixture (50 nM each hsa-miR-21-5p, hsa-miR-30a-3p and hsa-miR-93-5p) and antibodies through the 3-plex capture probe-loaded microbead field. (E) Shown are the percentages and box plots of mean fluorescence intensities of positive microbeads over time for the miRNA mixture diffusion. The contour plots, bar charts and box plots are exemplary for a representative experiment. All experiments were performed in triplicates. Circles indicate what has been added (blue = capture probe-loaded microbeads; red = matching miRNA; unfilled = non-matching miRNA). CP = capture probe, MB = microbead.

The miRNA binding to the capture probe-loaded microbeads could be observed in a time-dependent manner. The spatial distribution and the intensity of the fluorescence signals increased over time, due to successively exposed miRNAs to the capture probe-loaded microbeads. In this approach the capture probe-loaded microbeads in front of the chamber were first occupied and subsequently saturated with the miRNAs and non-bonded miRNA molecules diffused further and could therefore subsequently bind to the microbeads with free binding sites.

The same procedure was carried out with the diffusion approach II (miRNA diffusion in an antibody saturated environment). The difference was in the experimental setup. In contrast to the approach mentioned above, the antibodies were placed in the chamber together with the capture probe-loaded microbeads and only the miRNAs diffuse. With regard to specificity, a higher false positive rate of 7.7 % was found than for the diffusion approach I (3.3 %).

### Multiplex detection of miRNAs

In order to detect several miRNAs from a single sample (multiplex), we also tested the mixture of all three miRNAs (equal amounts of miRNAs) with both diffusion approaches. Co-diffusion of the miRNA mixture and antibodies against the capture probe-loaded microbeads (3-plex) is shown in Fig. 4D. Each miRNA of the mixture could be specifically detected in a time-dependent manner. Furthermore, diffusion kinetics were derived from the fluorescence data of the contour plots. These data are shown exemplar<u>il</u>y for the miRNA mixture in Fig. 4E. First, the number of microbeads that became positive over the diffusion time was calculated and displayed as bar plots. All beads whose capture probes were already occupied with miRNA, i.e., which had a significantly higher fluorescence intensity than the PBS-T control, were considered as positive microbeads. From all those positive microbeads, we calculated the mean fluorescence intensities and displayed them for each time point as box plots. The number of microbeads that had successfully bound miRNA increased over time, and the same beads that initially bound only small amounts of miRNA bound more free miRNAs depending on time. This led ultimately to an increased fluorescence intensity over time, which could be fitted to a four-parameter log-logistic curve. Same results were observed for diffusion approach II.

For each diffusion approach (I and II), three independent experiments were carried out and respective diffusion kinetics (see Fig. 4E) were prepared. The mean diffusion kinetics (Fig. 5) were then obtained from the kinetics of the individual experiments. Figure 5A depicts the diffusion kinetics for the number of positive microbeads. The comparison of the kinetics showed that in diffusion approach I (upper panel) about 40 % of the total microbeads became positive over time, whereas in diffusion approach II (lower panel) after 24 h almost 100 % of the microchamber surface was covered with positive microbeads. However, the comparison of the mean fluorescence intensities of these positive microbeads did not show any major differences between the two diffusion approaches (Fig. 5B). Furthermore, in this 3-plex mixture, the capture probe-loaded microbeads had strong specificity to their respective miRNA. There was no fluorescence signal on those microbead populations where non-complementary capture probes were used for the miRNA detection (Fig. 5C-E).

**Figure 5.**
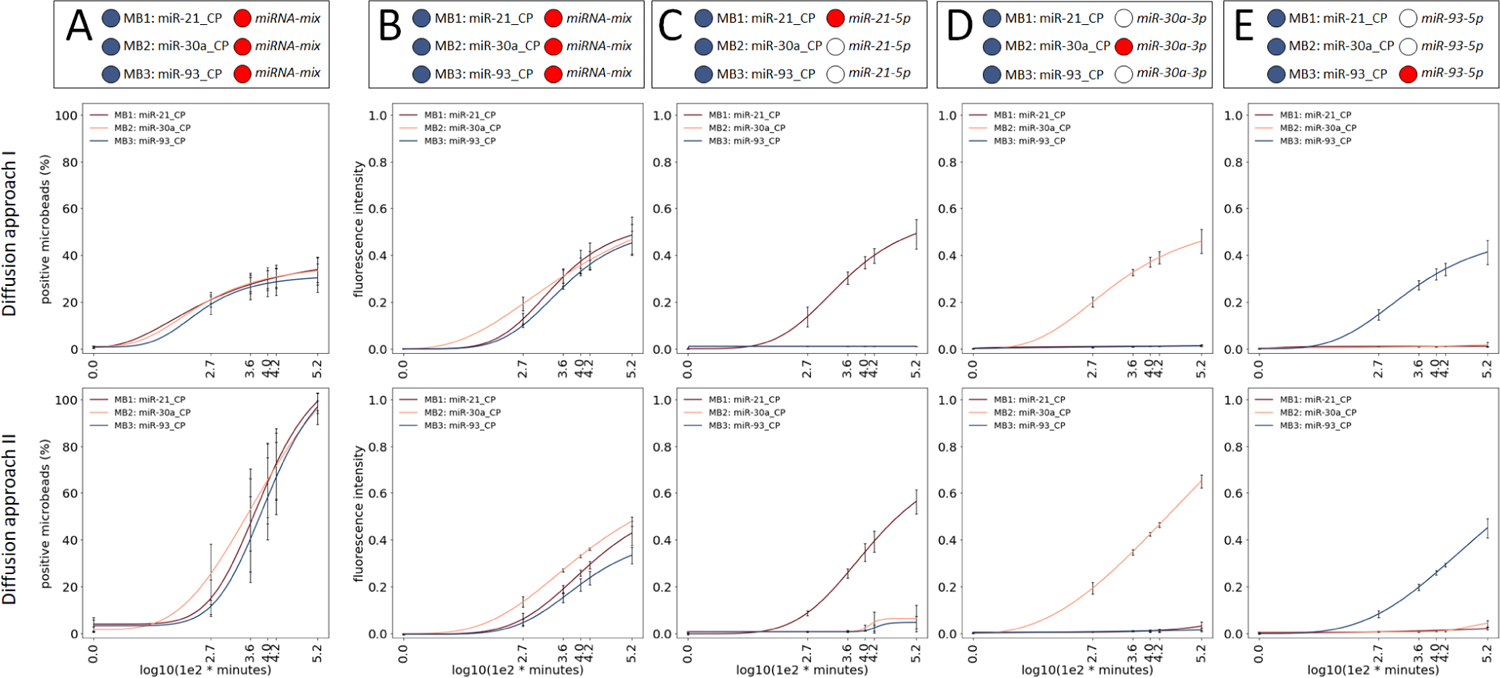
Diffusion kinetics for the multiplex antibody-based miRNA detection. Comparison of the kinetics of diffusion approach I and II regarding (A) number of positive microbeads in % for the miRNA mixture and mean fluorescence intensities of positive microbeads for (B) miRNA mixture, (C) hsa-miR-21-5p (50 nM), (D) hsa-miR-30a-3p (50nM) and (E) hsa-miR-93-5p (50 nM). Shown are mean ± standard deviation (n=3) for each diffusion approach. Circles indicate what has been added (blue = capture probe-loaded microbeads; red = matching miRNA; unfilled = non-matching miRNA). CP = capture probe, MB = microbead.

Different parameters can be determined from the diffusion kinetics. In order to compare the performance of both diffusion approaches, the second derivative was used. This gives us a statistic about the inflection point in the sigmoid curve and indicates when a signal in the assay starts to become positive. The second derivative was determined for the number of microbeads becoming positive as well as the change of fluorescence intensities over time.

### Sensitivity and detection limit of the microchamber diffusion assay

To evaluate the sensitivity of our diffusion-driven microchamber assay, we also investigated different miRNA concentrations ranging between 50 nM down to 0.05 nM as well as PBS-T control. The concentration-dependent miRNA detection was performed with both diffusion approaches (I and II) in the multiplex format.

### Diffusion approach I – co-diffusion of miRNAs and antibodies

Since we were able to show that no unspecific binding or cross hybridization of the miRNAs used in this assay occurred, the co-diffusion approach was performed with seven miRNA dilutions in 3-plex.

*Diffusion*: The contour plots for each miRNA were displayed individually and are exemplified for miR-21-5p in Figure 6A. As can be seen, we could specifically differentiate between various miRNA concentrations, which lead to different fluorescence intensities due to the respective miRNA dilution. With increasing miRNA concentration, the number of microbeads with successfully bound miRNAs increased (Fig. 6B). By using smaller amounts of miRNA, the percentage of positive microbeads also decreased, as the smaller number of miRNA molecules limits the radius of propagation in the chamber. At a miR-21-5p concentration of ≥ 5 nM, approximately 10-15 % of the total microbeads had already bound the miRNA and the two antibodies necessary for the detection of the hybrid within 5 minutes. After 40 minutes, about 30-40 % of microbeads were positive. The percentage number of positive microbeads in this assay has not increased further with increasing diffusion time. Only the fluorescence intensity changed depending on time and miRNA concentration (Fig. 6C). This means that microbeads which had already bound small amounts of miRNA and were therefore considered positive, bound more miRNA molecules over time. This lead to an increase in fluorescence intensity of one and the same beads.

**Figure 6.**
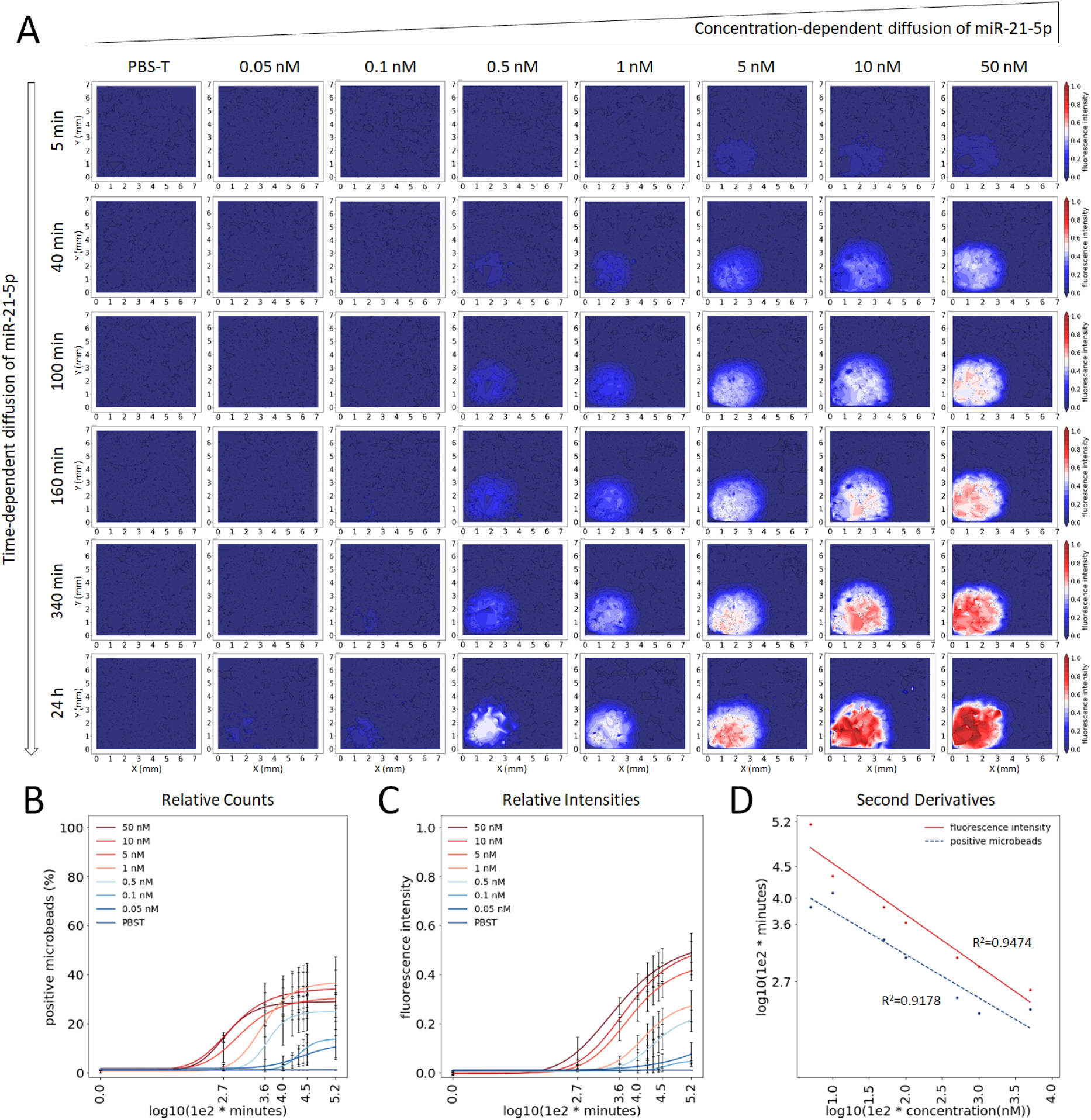
Diffusion approach I. (A) Contour plots of time- and concentration-dependent co-diffusion of miRNAs and antibodies exemplified by miR-21-5p. (B) Diffusion kinetics of the positive microbeads as a function of the miRNA concentration. Shown is the percentage number of microbeads for miR-21-5p as mean ± standard deviation (n=3). Microbeads with fluorescence intensity above the buffer control (mean + 3σ) are considered as positive. (C) Diffusion kinetics of miR-21-5p concentration-dependent mean fluorescence intensities. (D) The second derivative maximum of the diffusion time in minutes for the number of positive microbeads and the mean fluorescence intensities are plotted against the miRNA concentration. The labeling of the x- and y-axis are displayed as a logarithm of the concentration in nM, and the time point in minutes multiplied by 1e2, respectively.

In order to obtain a parameter for the sensitivity of this diffusion approach, the second derivative was derived from the concentration- and time-dependent fitted kinetics of the number of positive microbeads and the mean fluorescence intensities. The maximum of the respective second derivative (*SDM*) was plotted as a function of the miRNA concentration (Fig. 6D). In this case, the *SDM* referred to the time point in the curve at which the slope is maximum and the curvature is zero. With a concentration of 10-50 nM miR-21-5p, positive microbeads with an average fluorescence signal above the threshold value could be detected already after 1.5 minutes. The use of smaller amounts of miRNA extended the time to signal (5 nM after about 3 min, 1 nM after 10 min, 0.5 nM after 20 min, 0.1 nM and 0.05 nM after more than 100 min). The *SDM* of the mean fluorescence intensities of all positive microbeads was for 50 nM at about 4 min, 10 nM at 8 min, 5 nM at 12 min, 1 nM at 40 min, 0.5 at 70 min, 0.1 at 220 min and 0.05 at 1,400 min.

*Concentration-response curves*: Furthermore, we derived concentration-response curves from the data by plotting the miRNA concentration against the fluorescence intensities for each time point and calculated the *EC*_50_ values (Fig. S1-S3). The *EC*_50_ refers to the concentration, at which in the sigmoid curve the half maximal response is achieved. An improvement of the *EC*_50_ depending on the time can be observed for miR-21-5p (Fig. S1 A,C), miR-30a-3p (Fig. S2 A,C) and miR-93-5p (Fig. S3 A,C). But, after a diffusion time of 100 min no further improvement of the *EC*_50_ can be achieved (Fig. S1-S3 C). By calculating the *SDM* from the concentration-response curves, we see that, with the exception of miR-30a-3p, no higher sensitivity could be achieved due to continuing diffusion time (Fig. S1-S3 C).

### Diffusion approach II – miRNA diffusion in an antibody-saturated environment

*Diffusion kinetics*: The contour plots show that with diffusion approach II, in which the antibodies are already present in the chamber, a concentration- and time-dependent miRNA detection like approach I was possible (Fig. 7A). However, independent of the miRNA concentration used, no microbeads had bound miR-21-5p after 5 minutes diffusion time (Fig. 7B). With concentrations ≥ 5 nM, 30-40 % of the microbeads were positive after about 40 min. As the diffusion time increased, the percentage of positive microbeads increased even further up to 90 – 100 %, depending on the concentration. Within 24 hours, the entire chamber was completed with positive microbeads, at least at the higher miRNA concentrations. Lower miR-21-5p concentrations such as 1 or 0.5 nM reached a maximum of 60 % of the microbeads and with less than 0.5 nM detection is unlikely. The fluorescence intensities of positive microbeads increased over time in a concentration-dependent manner (Fig. 7C). We also derived the *SDM* from the fitted curves of Figure 7B and C. In contrast to diffusion approach I, with high miR-21-5p concentrations (10 and 50 nM) positive microbeads could only be detected after 25 and 20 minutes respectively (Fig. 7D). The time to signal increased by using lower miRNA concentrations (e.g. for 1 nM after 30 min, for 0.5 nM after 50 min, 0.1 nM and 0.05 nM after 130 min and 1,400 min, respectively). The *SDM* of the mean fluorescence intensities of all positive microbeads was for 50 nM at 12 min, 10 nM at 15 min, 5 nM at 25 min, 1 nM at 75 min, 0.5 at 110 min, 0.1 at 140 min and 0.05 at 1,400 min.

**Figure 7.**
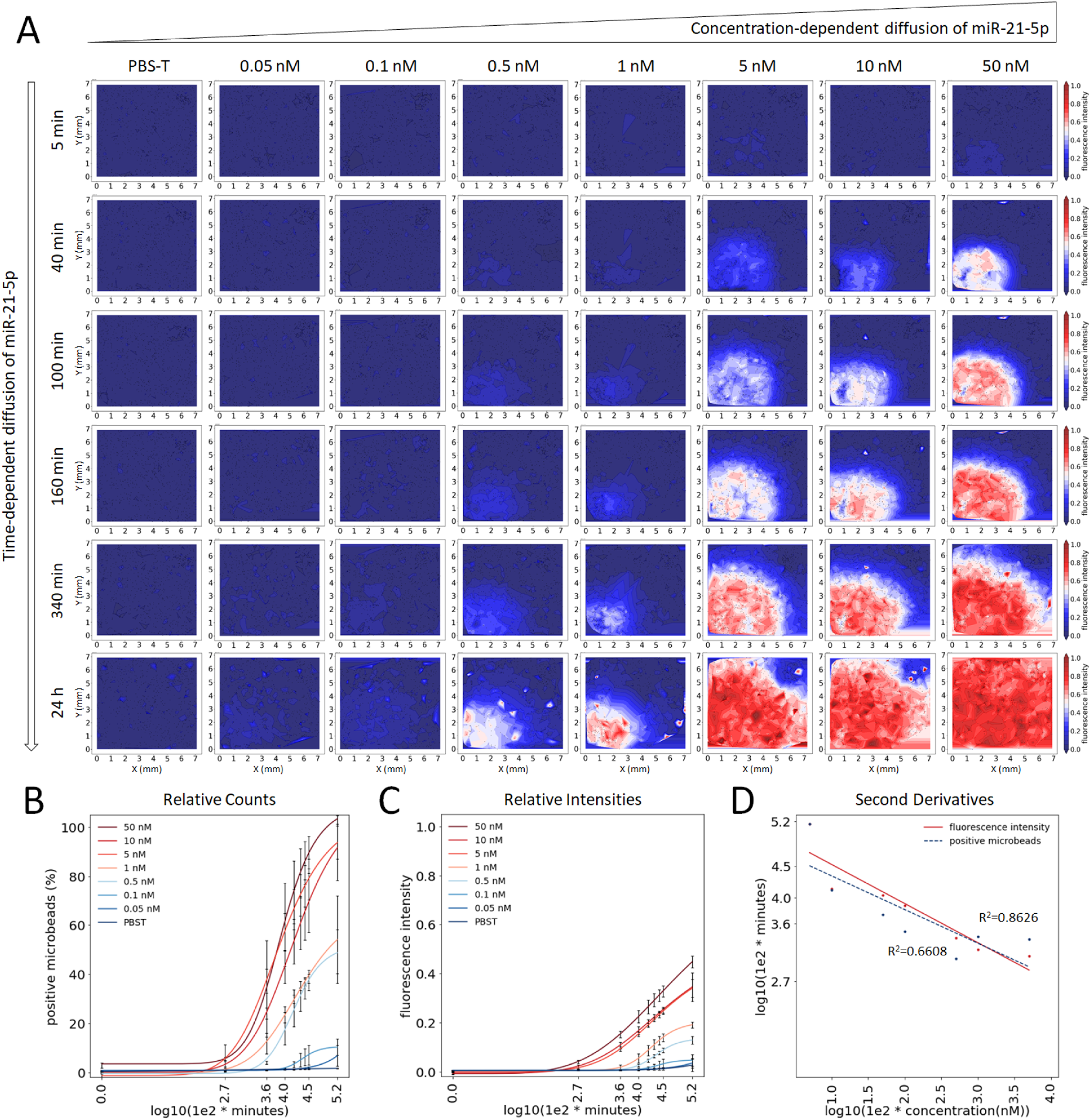
Diffusion approach II. (A) Contour plots of time- and concentration-dependent diffusion of miRNAs in an antibody saturated environment exemplified by miRNA-21. (B) Diffusion kinetics of the positive microbeads as a function of the miRNA concentration. Shown is the percentage number of microbeads for miR-21-5p as mean ± standard deviation (n=3). Microbeads with fluorescence intensity above the buffer control (mean + 3σ) are considered as positive. (C) Diffusion kinetics of miR-21-5p concentration-dependent mean fluorescence intensities (n=3). (D) The second derivative maximum of the diffusion time in minutes for the number of positive microbeads and the mean fluorescence intensities are plotted against the miRNA concentration. The labeling of the x- and y-axis are displayed as a logarithm of the concentration in nM, and the time point in minutes multiplied by 1e2, respectively.

**Figure S1.**
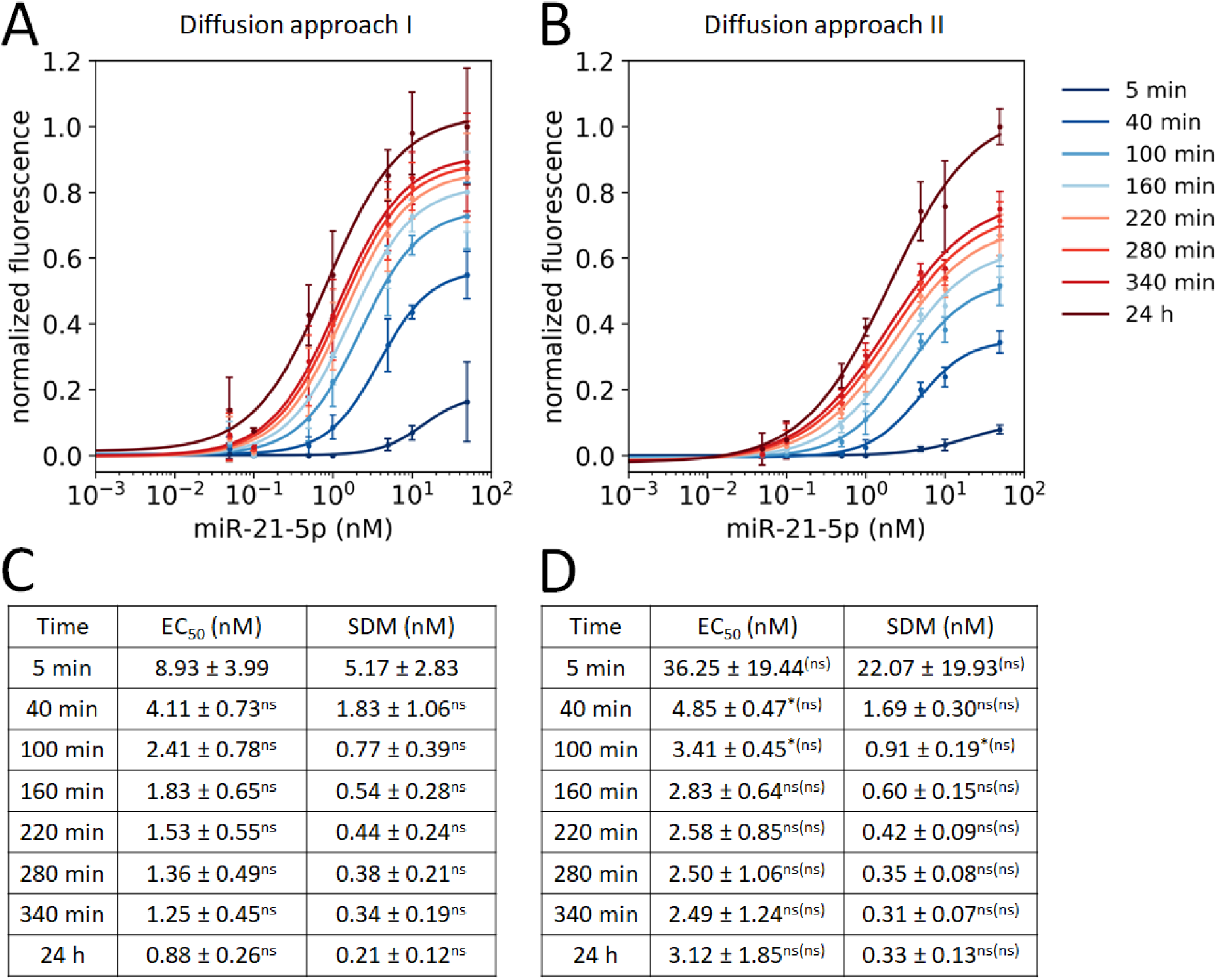
Hsa-miR-21-5p. Concentration-response curves (mean ± standard deviation) for diffusion approach (A) I and (B) II at different time points (n=3). The data were normalized to the maximum of the 24 hour measurement. Shown are EC_50_ and SDM values for hsa-miR-21-5p of diffusion approach (C) I and (D) II. The EC_50_ and SDM values were calculated for each individual experiment and displayed as mean ± standard deviation (n=3, two-sided t-test, ≤0.05*, ≤0.01**, ≤0.001***, nonsignificant (ns)). The significance levels between diffusion approach I and II are given in brackets. Significance levels between successive points in time within a diffusion assay are given without brackets.

**Figure S2.**
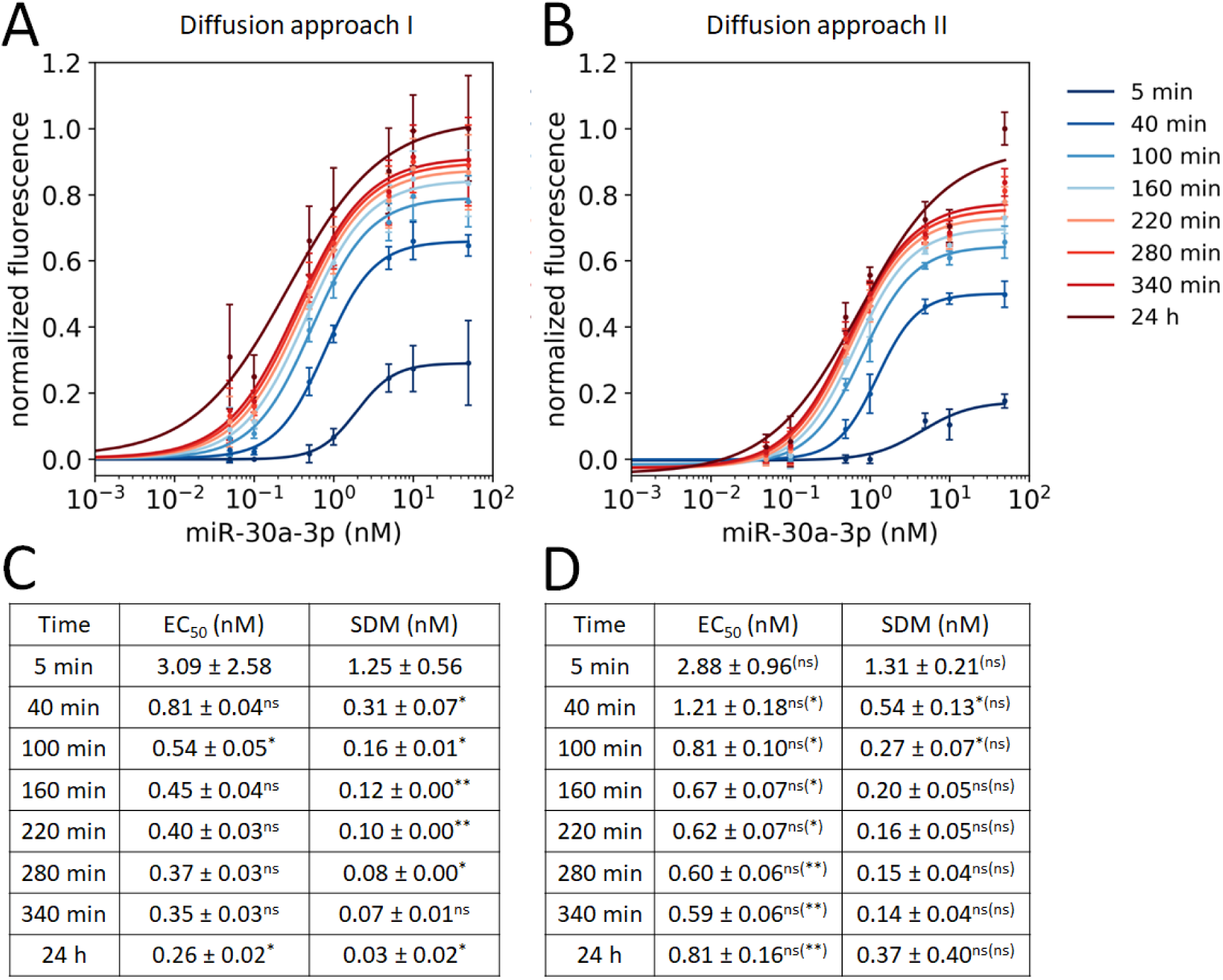
Hsa-miR-30a-3p. Concentration-response curves (mean ± standard deviation) for diffusion approach (A) I and (B) II at different time points (n=3). The data were normalized to the maximum of the 24 hour measurement. Shown are EC_50_ and SDM values for hsa-miR-30a-3p of diffusion approach (C) I and (D) II. The EC_50_ and SDM values were calculated for each individual experiment and displayed as mean ± standard deviation (n=3, two-sided t-test, ≤0.05*, ≤0.01**, ≤0.001***, nonsignificant (ns)). The significance levels between diffusion approach I and II are given in brackets. Significance levels between successive points in time within a diffusion assay are given without brackets.

**Figure S3.**
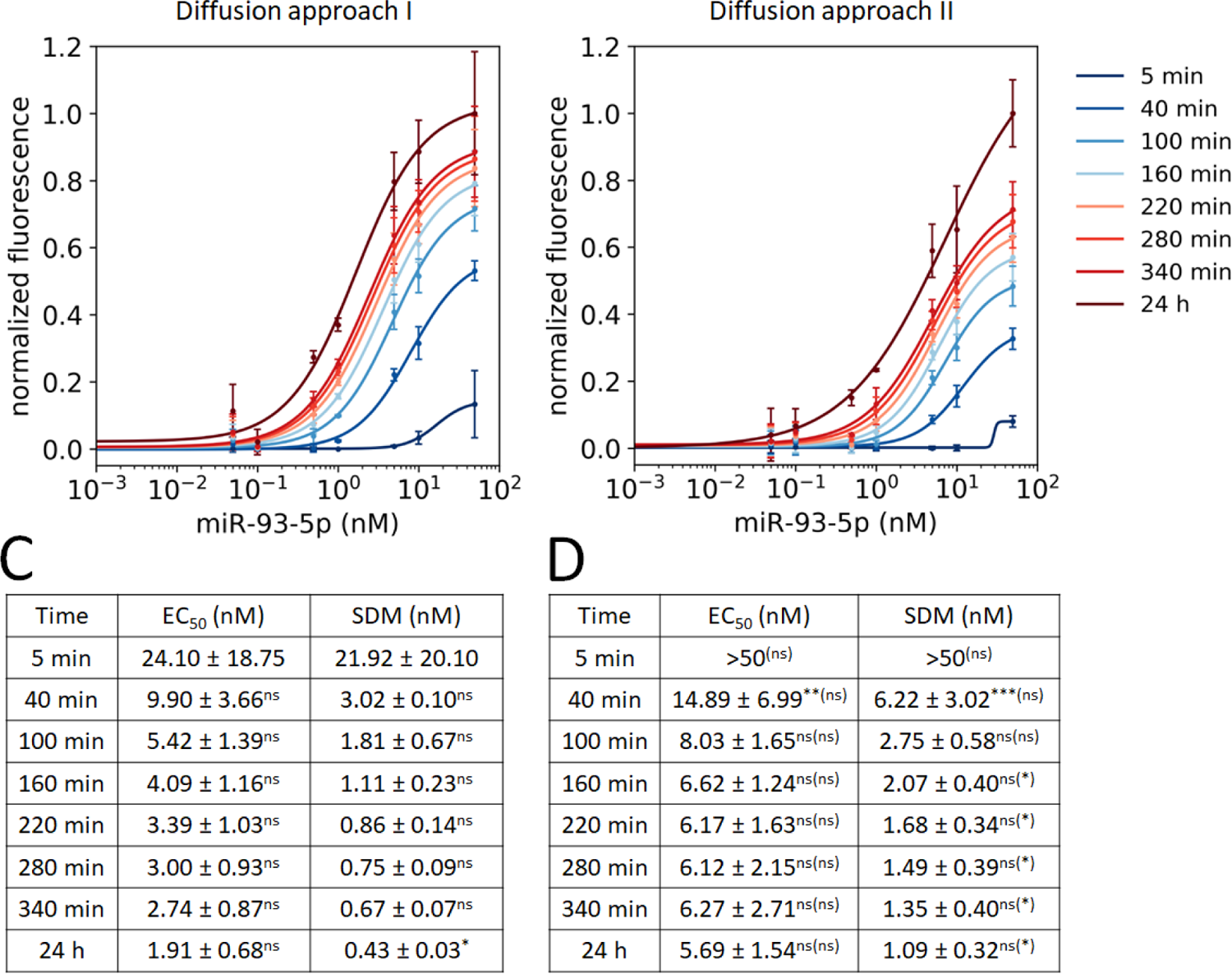
Hsa-miR-93-5p. Concentration-response curves (mean ± standard deviation) for diffusion approach (A) I and (B) II at different time points (n=3). The data were normalized to the maximum of the 24 hour measurement. Shown are EC_50_ and SDM values for hsa-miR-93-5p of diffusion approach (C) I and (D) II. The EC_50_ and SDM values were calculated for each individual experiment and displayed as mean ± standard deviation (n=3, two-sided t-test, ≤0.05*, ≤0.01**, ≤0.001***, nonsignificant (ns)). The significance levels between diffusion approach I and II are given in brackets. Significance levels between successive points in time within a diffusion assay are given without brackets.

**Figure S4.**
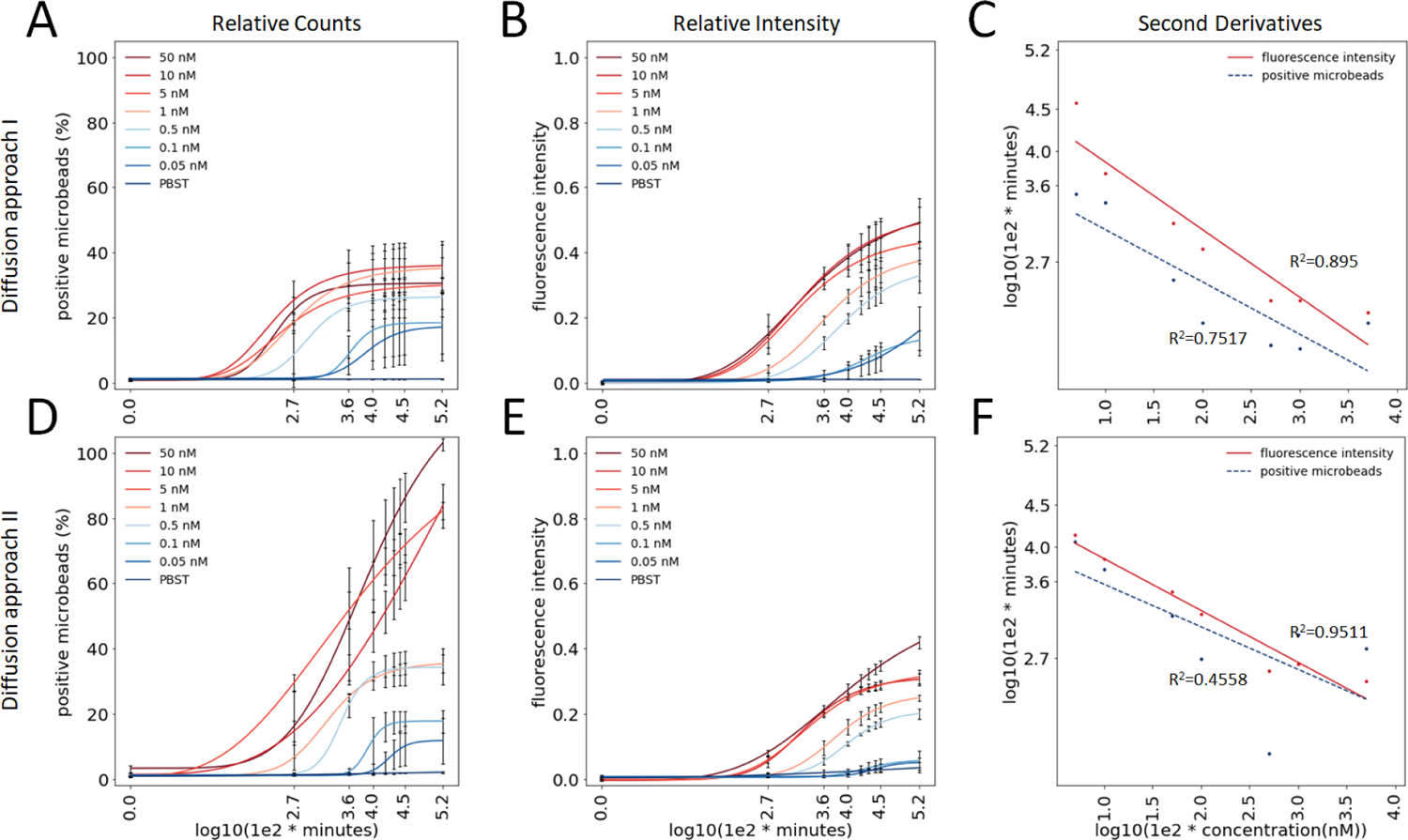
Comparison of the diffusion kinetics of hsa-miR-30a-3p for diffusion approach I and II. (A/D) Percentage change in the number of positive microbeads as a function of the miRNA concentration. (B/E) Change of mean fluorescence intensities of positive microbeads at different miRNA concentrations. Shown are mean ± standard deviation of n=3 each. (C/F) The second derivative maximum of the diffusion time in minutes for the number of positive microbeads and the mean fluorescence intensities are plotted against the miRNA concentration.

**Figure S5.**
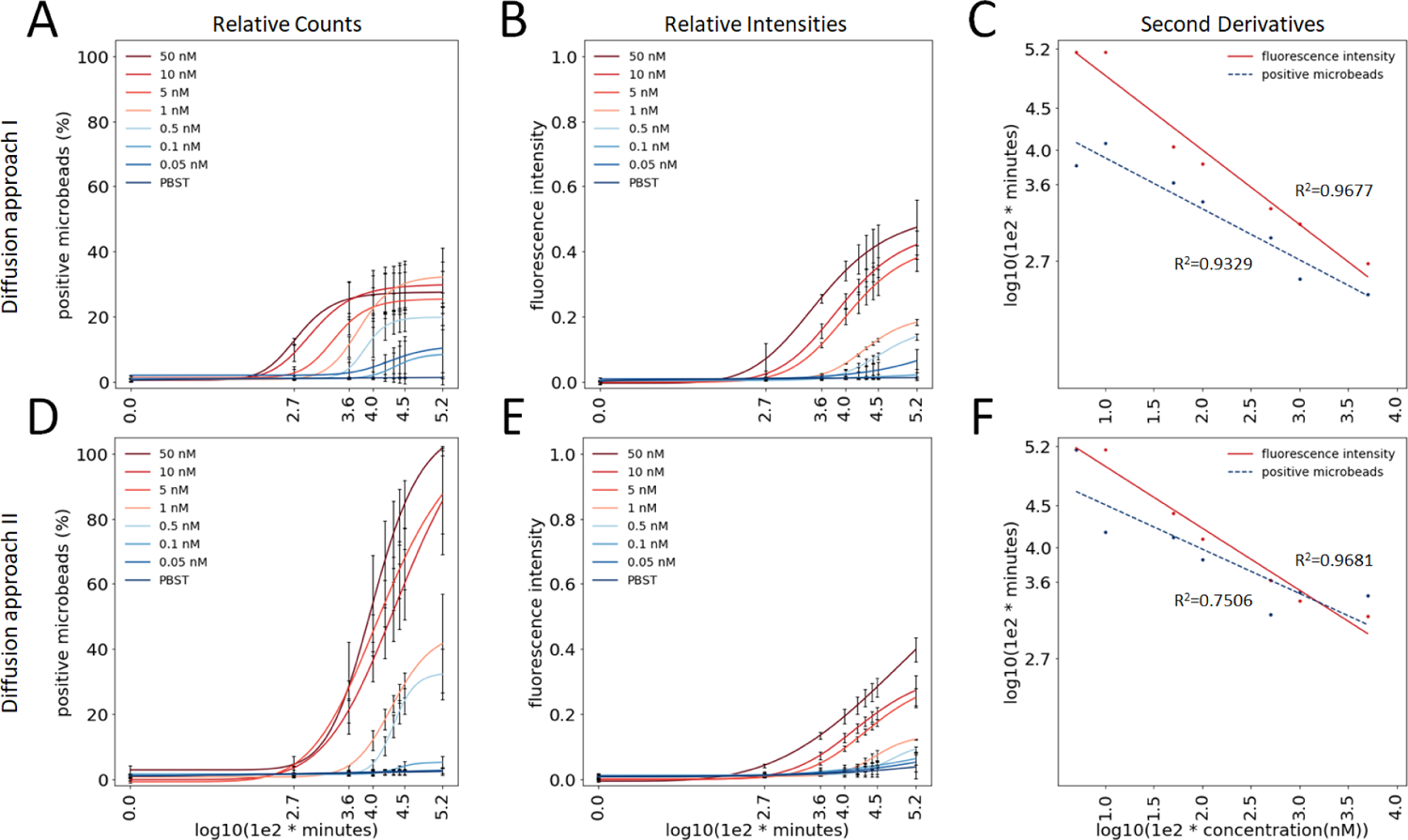
Comparison of the diffusion kinetics of hsa-miR-93-5p for diffusion approach I and II. (A/D) Percentage change in the number of positive microbeads as a function of the miRNA concentration. (B/E) Change of mean fluorescence intensities of positive microbeads at different miRNA concentrations. Shown are mean ± standard deviation of n=3 each. (C/F) The second derivative maximum of the diffusion time in minutes for the number of positive microbeads and the mean fluorescence intensities are plotted against the miRNA concentration.

*Concentration-response curves*: We also derived concentration-response curves for miR-21-5p (Fig. S1 B,D), miR-30a-3p (Fig. S2 B,D) and miR-93-5p (Fig. S3 B,D) from the data of diffusion approach II and calculated *EC*_50_ values from the fitted curves. After at least one hour diffusion time (between 40 and 100 minutes time points), there was no further change in the *EC*_50_ values (Fig. S1-S3 D), i.e. with regard to this parameter, which is commonly used to determine affinity and thus sensitivity, a further incubation time did not lead to an improvement, which suggests that the miRNA detection had already been completed. Comparison of the time-dependent *SDM* values revealed the same (Fig. S1-S3 D).

### Comparison of diffusion approach I and II

When comparing the two diffusion approaches, it can be seen that each has advantages and disadvantages with regard to the parameters listed in Table 3.

**Table 3.** Comparison of both diffusion approaches. A plus sign indicates a positive evaluation and defines the ranking.

|  | **Diffusion approach I**<br>co-diffusion of<br>miRNA and antibodies | **Diffusion approach II**<br>miRNA diffusion in an<br>antibody saturated-<br>environment |
| --- | --- | --- |
| Handling | ++ | +++ |
| Specificity | +++ | ++ |
| Multiplexing | +++ | +++ |
| *Signal-to-noise ratio:* |  |  |
| Number of positive microbeads | + | +++ |
| Mean fluorescence intensity | +++ | ++ |
| *Sensitivity:* |  |  |
| Time to signal | +++ | ++ |
| EC<sub>50</sub> | +++ | ++ |
| SDM | +++ | ++ |

*Handling, specificity and multiplexing*: The three investigated miRNAs could be detected specifically and in multiplex with both approaches. A rather better assay specificity (96.7 % *vs.* 92.3 %) could be achieved with the first approach. The diffusion approach II offers a rather easier handling compared to I, because the antibodies do not have to be added to each individual miRNA dilution. By adding the antibodies together with the capture probe-loaded microbeads in the chamber, assay errors might be reduced.

*Signal-to-noise ratio*: We observed detection differences depending on the experimental approach regarding the number of microbeads becoming positive over time and the overall fluorescence intensities. With the same amount of miRNA as in diffusion approach I, a larger detectable propagation radius was achieved in diffusion approach II (Fig. 6A, Fig. 7A). With diffusion approach I only 30-40% of all microbeads become positive for miR-21-5p (Fig. 6B), miR-30a-3p (Fig. S4 A) and miR-93-5p (Fig. S5 A), whereas with approach II for all three miRNAs 90-100 % (Fig.7B, Fig. S4 D, Fig. S5 D) are achieved. Nevertheless, higher fluorescence intensities can be achieved with the first approach for all three miRNAs than with the second (Fig. 6B, Fig. 7B, Fig. S4 B,E, Fig. S5 B,E), which can be confirmed by better signal-to-noise ratios (Table S3).

*Sensitivity*: Although diffusion approach I seems to be more sensitive than approach II with regard to *EC*_50_ and *SDM* values, no significant differences can be determined, with the exception of miR-30a-3p for *EC*_50_ or miR-93-5p for *SDM* (Fig. S1-S3 C,D). The time to signal with respect to the number of positive microbeads and relative fluorescence intensities is faster with diffusion approach I than with diffusion approach II. Smaller amounts of miRNA can be detected earlier with approach I.

Since all conditions except the presence of the antibodies (co-diffusion or already present) were the same, this seems to be the limiting factor in approach I. Limiting at least in terms of the number of microbeads that became positive over time. This could be due to different diffusion rates of miRNA and antibodies. They differ in mass by the factor of about 21 [33], because a miRNA weighs about 7 kDa and an IgG antibody approximately 150 kDa. The miRNAs were able to spread over the entire microchamber within 24 h by diffusion. This could be demonstrated by the approach II. In approach I, the antibodies, which are ultimately the signal generators, remained behind the miRNAs, i.e. the number of positive microbeads (in the maximum 40 %) refers to the propagation limit in which miRNA and antibodies still overlap. MiRNAs diffused further to the end of the chamber. They reached this end after about 24 h. The antibodies were lagging behind due to their possible inertia, i.e. they might take longer than 24 h to diffuse further and bind free DNA:miRNA hybrids. The number of positive microbeads depends equally on how many miRNA molecules have bound and how well the DNA:miRNA hybrid is recognized by the primary and secondary antibody. A miRNA bound solely to the complementary capture probes of the microbead cannot be detected by *VideoScan*. The entire microbead complex with probe + miRNA + primary antibody + secondary antibody is always required for the detection.

Another aspect is the gradient in the chamber itself. This applies to both approaches and to each dilution stage. There is always a gradient from a higher miRNA concentration at the entrance of the chamber to a lower concentration at the opposite end. This is due 1) to the fact that miRNA molecules bind to probes at the very beginning of the process and thus can diffuse less and less further until at the end all molecules are completely bound or an excess remains because all probes on the microbeads are already occupied. The latter could occur at too high concentrations. In the case of very small amounts of miRNA, the molecules are used up more quickly, i.e. bound to probes, and diffusion ends earlier. 2) It is due to the experimental dilution factor itself. Liquid in a volume of 7 µL is presented in the chamber and the miRNA (with or without antibodies, depending on the approach) is dissolved in a volume of 2 µL. Mixing of both volumes results in gradual dilution.

Although the sensitivity of the presented miRNA detection method cannot compete with quantitative PCR, there are advantages in hybridizing miRNA with capture probe-loaded microbeads which are reached stepwise over time (diffusion assays), compared to those, which are simultaneously present (bulk mixed, reference assays) (Fig. S6-S8). In the reaction vessel, the hybridization of the miRNAs to the capture probes is non-directed and in the diffusion microchamber a directed propagation can be ensured. The diffusion chamber offers the possibility to count the microbeads, namely microbeads that become positive depending on time and miRNA concentration. It can also be clearly defined at which location a signal has to be emitted and at which not. False positive signals can therefore be excluded with the diffusion approach. This cannot be achieved in the reaction vessel, because all microbeads are mixed simultaneously and evenly.

It was shown that real-time DNA microarrays have several advantages over end-point measurements. Real-time systems have enhanced detection dynamic range and they are less dependent on probe saturation in the capturing spots and processing steps such as washing artifacts and microarray spot-to-spot variations [34].

The diffusion assay is suitable for multiplex detection of several miRNAs from one sample. Due to the tiny reaction cavity of the chamber, it is sample-saving and therefore applicable for small sample material. The presumed advantage is, that the miRNAs are successively exposed to the capture probe-loaded microbeads and that in this approach the capture probe-loaded microbeads in front are first saturated with the target molecules and non-bonded miRNAs diffuse further and can therefore subsequently bind to the microbeads with free binding sites further inside the chamber.

Briefly, the microbeads can be digitized into two groups: Firstly positive microbeads to which the miRNA has specifically bound (positive, 1) and secondly microbeads to which no miRNAs have bound within the elapsed time (negative, 0). Each microbead represents a measuring point that can be used to quantify the fluorescence intensity of the miRNAs (continuous variable) on the microbead. Since a certain number of miRNAs can bind specifically to each microbead until saturation, the signals increase over time. However, the signal increase is only noticeable at the microbeads, where the analyzed solution with the miRNA has already diffused. The other microbeads therefore have a negative signal. The diffusion of miRNAs is similar to a random walk.

So far, we are not using the available information of all the data types (discrete state (positive, negative) and continuous quantity) for the quantification of miRNAs. The diffusion assay provides the prerequisite for quantification. Quantification of miRNA is generally important because its expression (high/low) plays a role in many diseases, e.g. in the development of disease or its progress monitoring. The miRNAs could be quantified in our assay e.g. by the spreading radius, i.e. by the time of microbeads becoming positive. This radius depends on the miRNA concentration (for this purpose, an extra software tool would have to be developed).

Nevertheless, some limitations should be addressed. Diffusion is a slow process and therefore time-consuming. This fact, combined with the complex sample preparation, makes the application in a diagnostic laboratory currently challenging. Therefore, we intend to integrate this approach into a microfluidic platform to optimize handling and incubation time. For the test to be applicable, its sensitivity must be improved for routine diagnostics. Although it is possible to detect miRNAs in the nanomolar range, these concentrations are nevertheless above the picomolar levels found in body fluids such as plasma or serum. To solve this problem, we will try to validate buffer and temperature conditions, change the chamber geometry (e.g. microfluidics), use dendrimers or more intensely labeled antibodies and in addition antibodies against selected miRNAs, which can detect these miRNAs with a high sensitivity. We aim to gradually increase the multiplex level of the assay to allow the determination of as many miRNAs as possible from a single sample.

## Conclusion

Our microbead-based microchamber diffusion assay is suitable for the antibody-based detection of miRNAs. By directly detecting a resulting DNA:miRNA hybrid with an anti-DNA:RNA hybrid antibody, no labeling of the miRNA is necessary, which renders the assay universal and suitable for the detection of any relevant miRNA. The specificity of the assay can be guaranteed by using miRNA-specific complementary probes and the anti-DNA:RNA hybrid antibody only recognizes perfectly matching hybrids [15, 26, 35]. Furthermore, this assay allows the isothermal detection of miRNAs with both primary and secondary antibodies in a single reaction step without any amplification processes. To the best of our knowledge, this is the first assay that combines a directed diffusion of the analyte solution with a planar multiplex microbead array and antibody-based detection of miRNA and real-time monitoring.

The basic idea of the microchamber diffusion assay can be useful for other applications, e.g. for cell culture experiments in which specific target molecules diffuse past cells, receptor interaction studies where activators or inhibitors diffuse in a directed manner or generally for the detection of other nucleic acids or proteins.

## Acknowledgments

We thank Philipp Müller, Alexander Böhm and Jörg Nitschke (BTU Cottbus-Senftenberg) for their contributions to the development of the assay.

## Supplemental Information

**Table S1.** *Thermodynamic parameters of miRNAs used in this study. Duplex formation of miRNA and DNA oligonucleotides in solution were calculated by nearest-neighbor analysis with the unified parameters in 1 M NaCl* [36].

| miRNA | melting temperature<br>(calculated)<br>TM [°C] | delta Gibbs free energy<br>(calculated)<br>$\Delta G^0$ [kJ/mol] | delta enthalpy<br>(calculated)<br>$\Delta H^0$ [kJ/mol] | delta entropy<br>(calculated)<br>$\Delta S^0$ [J/(mol·K)] |
| --- | --- | --- | --- | --- |
| miR-21-5p | 50.05 | -94.98 | -721.74 | -2,020.87 |
| miR-30a-3p | 51.05 | -102.93 | -835.96 | -2,363.12 |
| miR-93-5p | 57.81 | -119.24 | -842.66 | -2,332.16 |

**Table S2.** *Logarithmic time and concentration*.

| time in minutes | $\log_{10}(1e2*\text{minutes})$ | concentration in nM | $\log_{10}(1e2*\text{concentration(nM)})$ |
| --- | --- | --- | --- |
| 5 | 2.7 | 0.00 | — |
| 40 | 3.6 | 0.05 | 0.7 |
| 100 | 4.0 | 0.1 | 1.0 |
| 160 | 4.2 | 0.5 | 1.7 |
| 220 | 4.3 | 1 | 2.0 |
| 280 | 4.4 | 5 | 2.7 |
| 340 | 4.4 | 10 | 3.0 |
| 1,440 | 5.2 | 50 | 3.7 |

**Table S3.**
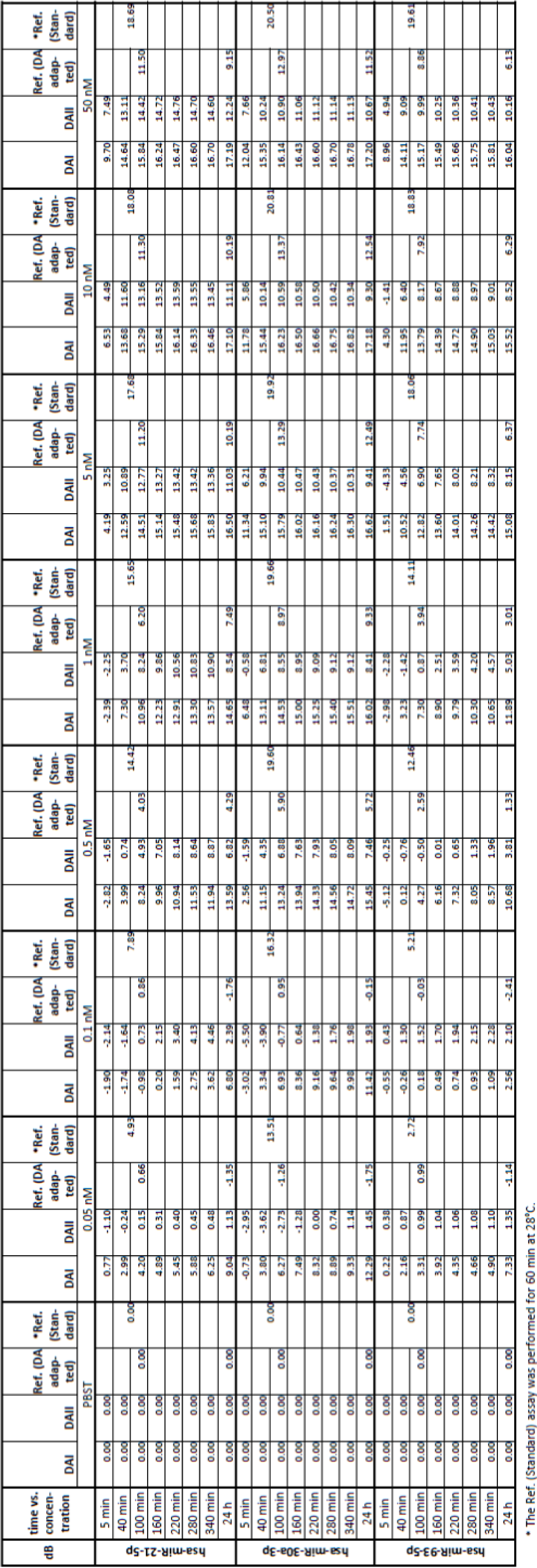
*Signal-to-noise ratios. The signal-to-noise ratios (SNR) were calculated for the diffusion assay I (DAI), the diffusion assay II (DAII), the reference assay adapted to the diffusion assay (Ref. DA adapted) and the standard reference assay (Ref. Standard) by the equation SNR = 10log <u>^signal^</u>, where signal is the fluorescence intensity of the tested miRNAs at different concentrations and time points, and noise the fluorescence intensity of the respective PBST control. Higher dB values indicate better signal-to-noise ratios*.

**Table S4.**
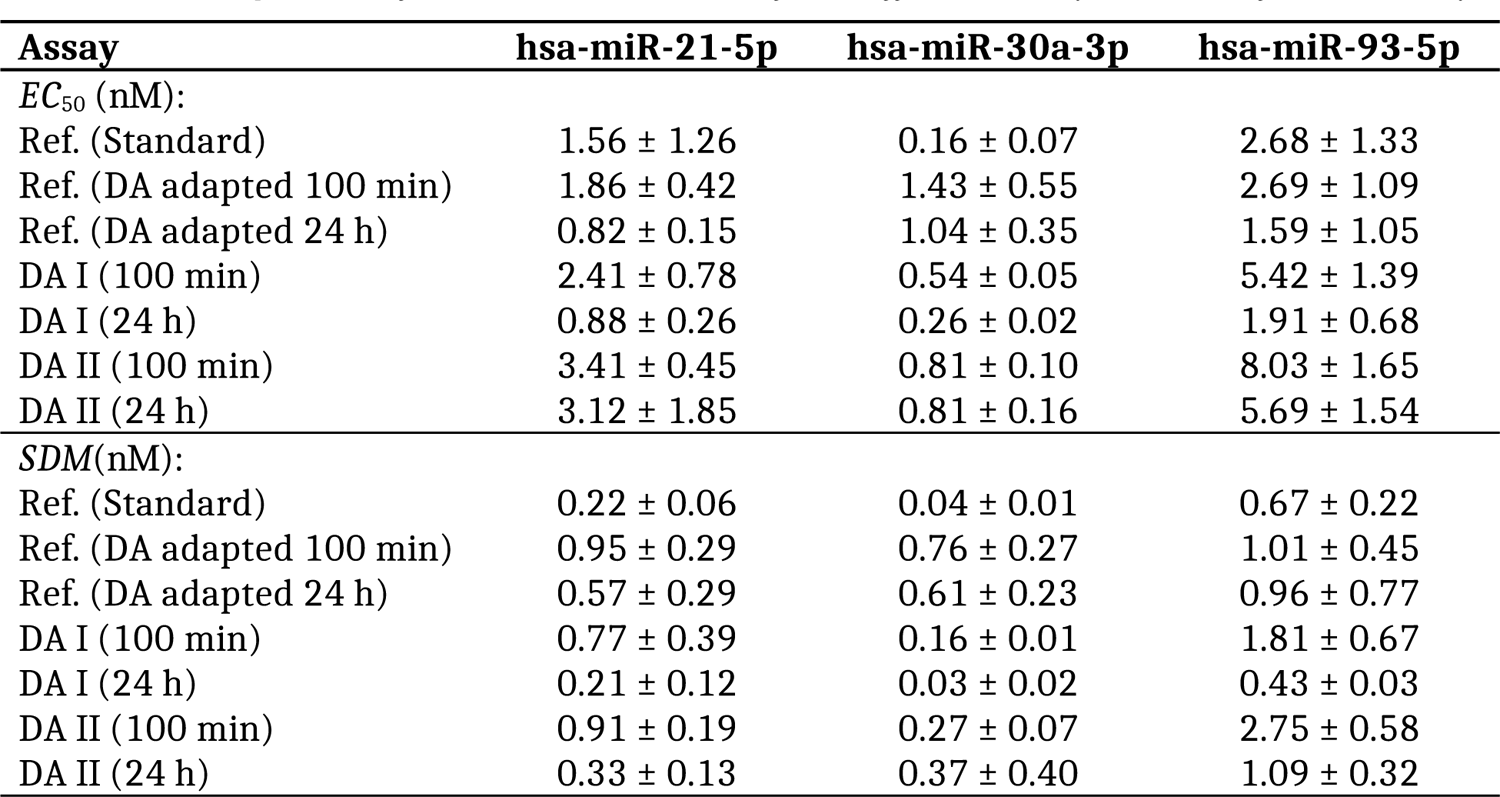
*Comparison of EC_50_ and SDM values of the diffusion assay and the reference assay*.

| Assay | hsa-miR-21-5p | hsa-miR-30a-3p | hsa-miR-93-5p |
| --- | --- | --- | --- |
| *EC<sub>50</sub>* (nM): |  |  |  |
| Ref. (Standard) | 1.56 ± 1.26 | 0.16 ± 0.07 | 2.68 ± 1.33 |
| Ref. (DA adapted 100 min) | 1.86 ± 0.42 | 1.43 ± 0.55 | 2.69 ± 1.09 |
| Ref. (DA adapted 24 h) | 0.82 ± 0.15 | 1.04 ± 0.35 | 1.59 ± 1.05 |
| DA I (100 min) | 2.41 ± 0.78 | 0.54 ± 0.05 | 5.42 ± 1.39 |
| DA I (24 h) | 0.88 ± 0.26 | 0.26 ± 0.02 | 1.91 ± 0.68 |
| DA II (100 min) | 3.41 ± 0.45 | 0.81 ± 0.10 | 8.03 ± 1.65 |
| DA II (24 h) | 3.12 ± 1.85 | 0.81 ± 0.16 | 5.69 ± 1.54 |
| *SDM*(nM): |  |  |  |
| Ref. (Standard) | 0.22 ± 0.06 | 0.04 ± 0.01 | 0.67 ± 0.22 |
| Ref. (DA adapted 100 min) | 0.95 ± 0.29 | 0.76 ± 0.27 | 1.01 ± 0.45 |
| Ref. (DA adapted 24 h) | 0.57 ± 0.29 | 0.61 ± 0.23 | 0.96 ± 0.77 |
| DA I (100 min) | 0.77 ± 0.39 | 0.16 ± 0.01 | 1.81 ± 0.67 |
| DA I (24 h) | 0.21 ± 0.12 | 0.03 ± 0.02 | 0.43 ± 0.03 |
| DA II (100 min) | 0.91 ± 0.19 | 0.27 ± 0.07 | 2.75 ± 0.58 |
| DA II (24 h) | 0.33 ± 0.13 | 0.37 ± 0.40 | 1.09 ± 0.32 |

## Characterization of DNA:miRNA hybrids

For the three miRNAs in this study, we also calculated the melting temperature and some other miRNA properties (at 37 °C) such as delta Gibbs free energy (Δ*G*^0^), delta enthalpy (Δ*H*^0^) and delta entropy (Δ*S*^0^), see Table S1. The Δ*G*^0^ of forming the DNA:miRNA hybrids was calculated using the nearest-neighbor method, and the nearest-neighbor parameters for DNA:RNA duplexes in 1 M NaCl buffer was from Sugimoto *et al.* [36]. The knowledge about the theoretical melting temperature was used during the assay design. In particular, an incubation temperature of 24 °C was chosen as a starting point with stable DNA:miRNA hybrids.

## Dose-response curve fitting

The logarithm conversion values for time and concentration are given in Table S2.

## Sensitivity and detection limit of the microchamber diffusion assay

Supporting information of this subsection include Fig. S1, Fig. S2, Fig. S3, Fig. S4, Fig. 5 and Table S3.

## Reference assay

### Standard

The miRNA hybridization and antibody binding was performed in a 200 µL reaction vessel. In this assay, the capture probe-loaded microbeads (3-plex; approx. 1,000 beads per population; see above) were added together with the respective miRNAs (serial dilutions as mentioned above), the anti-DNA:RNA hybrid antibody and the secondary antibody in the reaction vessel. The standard volume of the whole reaction mix was 50 µL. After 1 h incubation at 28°C with vigorous shaking (1,200 rpm), the microbeads were washed three times with PBS-T, transferred to a 96-well plate, and the fluorescence signals were analyzed with *VideoScan* (4 images per well).

## Adapted to the diffusion assay

Since the volume and concentration ratios between the standard assay and the microchamber diffusion assay differ, the standard assay was adapted. Here, the same volume and concentration ratios were used as in the diffusion microchamber. In order to achieve a sufficient mixing of the capture probe-loaded microbeads, miRNAs and antibodies in the 200 µL reaction vessel, only the total reaction volume was increased (2-fold) without changing the ratios to each other. The reaction mix was incubated at 24 °C and vigorous shaking (1,200 rpm). After 100 min and 24 h incubation time, 9 µL (total volume in the diffusion microchamber) were taken from the reaction mixture, washed three times with PBS-T, transferred to a 96-well plate, and the fluorescence signals were analyzed with *VideoScan* (4 images per well).

## Comparison of the microchamber diffusion assay with the reference assay

In both diffusion approaches, the hybridization of the miRNA takes place stepwise to the capture probe loaded microbeads, which are randomly ordered on the bottom surface of the chamber. In contrast, miRNA hybridization was performed in a reaction vessel in which all molecules are exposed to the microbeads simultaneously (Fig. S6 A).

**Figure S6.**
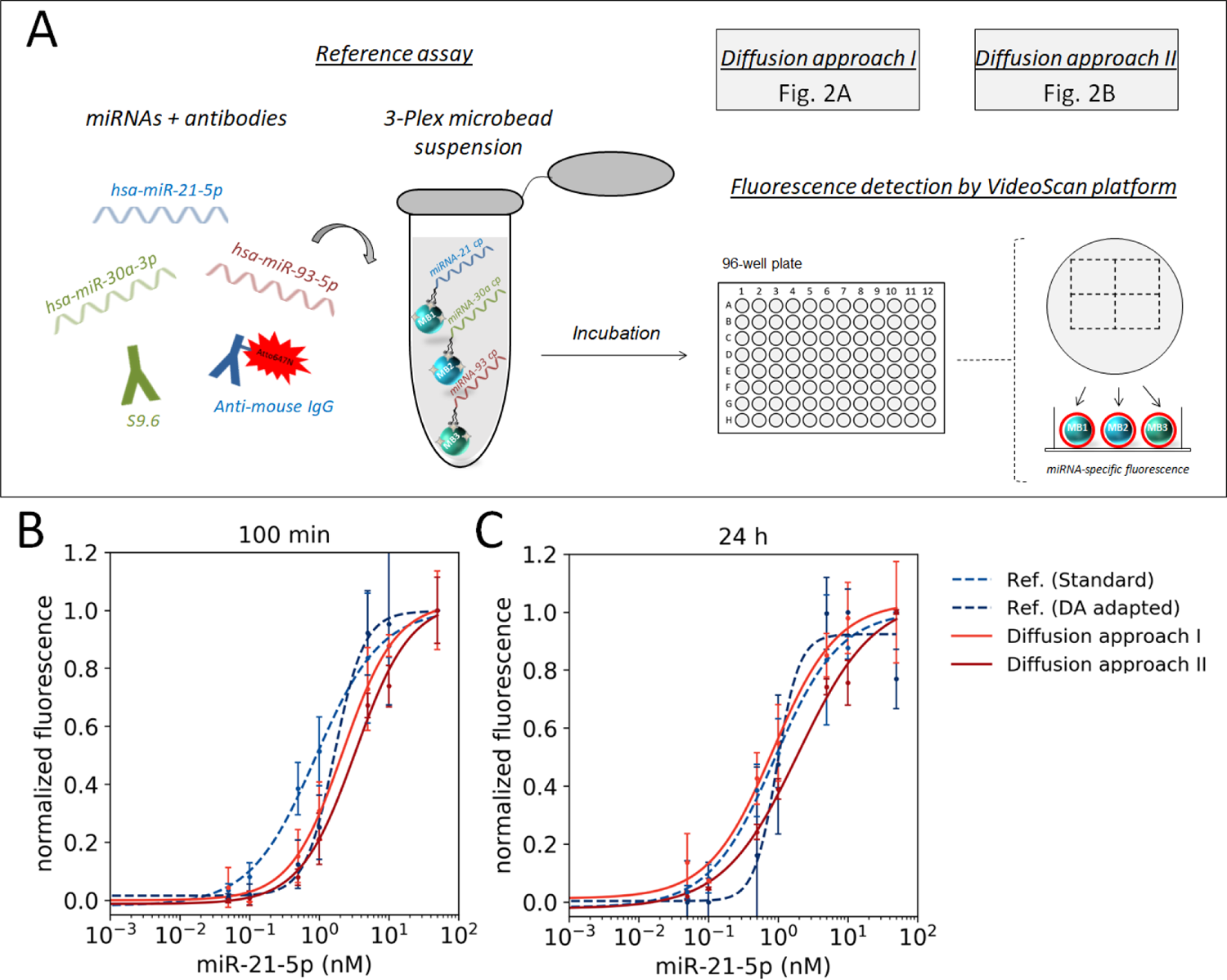
Comparison of the microchamber diffusion assay with the reference assay (A) Cartoon of the reference assay. The miRNA hybridization and antibody binding takes place simultaneously in one reaction vessel and the fluorescence is then measured in 96-well format by VideoScan. For the adapted reference assay, the volume and concentration ratios as well as the incubation times of the diffusion assay were used. Comparison of the concentration-response curves exemplified by miR-21-5p at timepoints (B) 100 min and (C) 24 h. The standard reference assay curve is the same in panel (B) and (C). Shown are mean ± standard deviation (n=3). Ref.(Standard) = standard reference assay. Ref.(DA adapted) = reference assay adapted to the diffusion assay.

**Figure S7.**
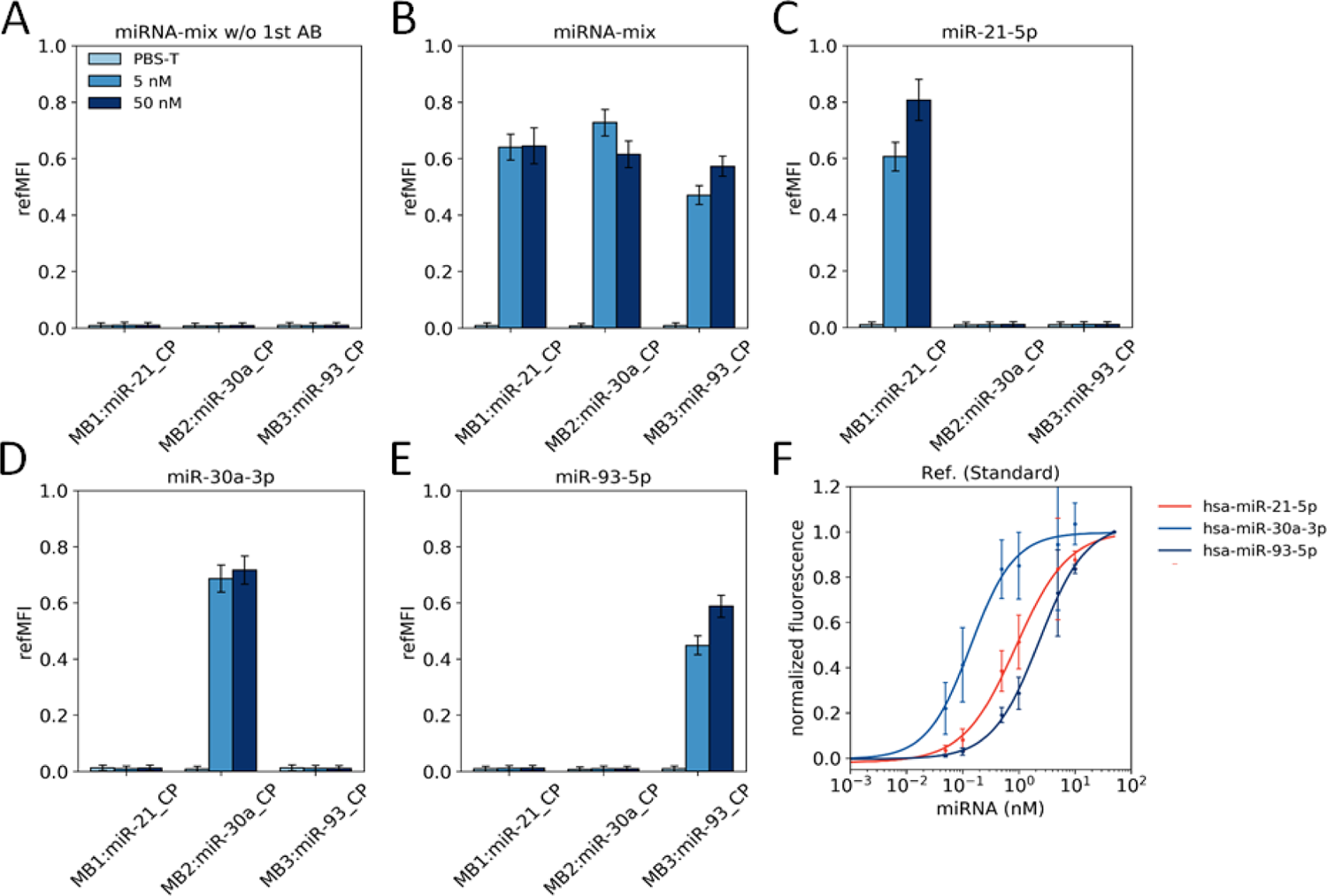
Bulk mixed endpoint measurement. The capture probe-loaded microbeads (3-plex), miRNAs (as mixture or individually), primary and secondary antibody were mixed and incubated for 1 h at 28 °C with vigorous shaking. Shown are multiplex detection of (A) miRNA mixture without primary antibody, (B) miRNA mixture, (C) hsa-miR-21-5p, (D) hsa-miR-30a-3p, (E) hsa-miR-93-5p and (F) concentration-response curves as mean ± standard deviation of n=3 experiments. Ref. (Standard) = reference assay standard method, MB = microbeads, CP = capture probe.

**Figure S8.**
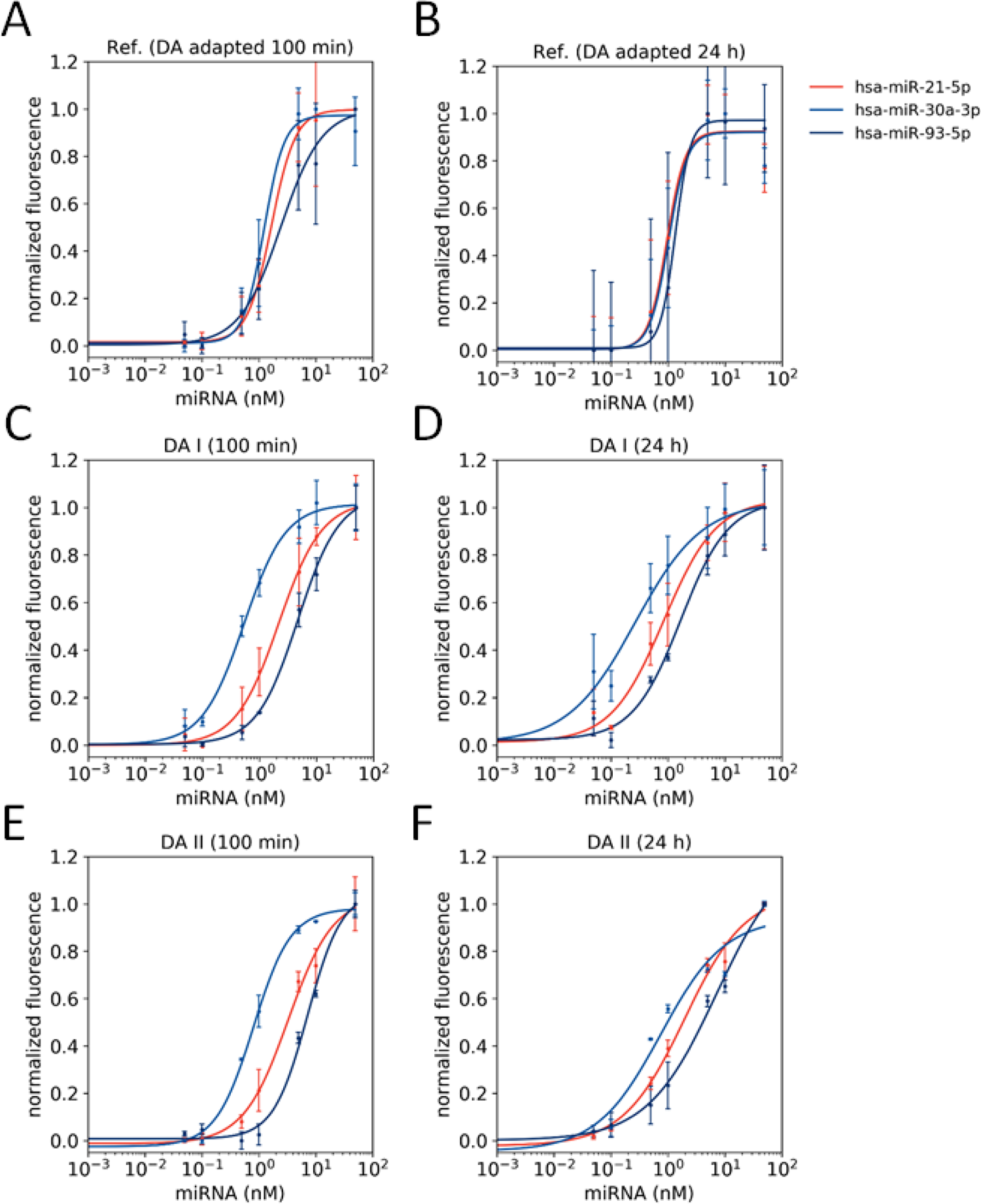
Comparison of the diffusion assay with the reference assay. Shown are concentration-response curves of (A) diffusion assay adapted reference assay at 100 min, (B) diffusion assay adapted reference assay at 24 h, (C) diffusion approach I at 100 min, (D) diffusion approach I at 24 h, (E) diffusion approach II at 100 min and (F) diffusion approach II at 24 h as mean ± standard deviation of n=3 experiments. Ref. (DA adapted) = reference assay adapted to the diffusion assay, DA I/ DA II = diffusion approach I/ II

In the standard reference assay, miRNA, both antibodies and the microbeads carrying capture probes are incubated in a defined reaction volume for a defined time and then the entire reaction mixture is measured by fluorescence microscopy in 96-well format. This reference assay has also been modified by adapting it to the diffusion assay in terms of volume and concentration ratios, temperature and incubation times. This is referred to hereinafter as the adapted reference assay.

The comparison of the concentration-response curves is exemplified for miR-21-5p after 100 min in Fig. S6 B and after 24 h in Fig. S6 C. The curve for the standard reference assay is the same in both panels, as the incubation time is 1 h and the temperature 28 °C. For the adapted reference assay, the diffusion assay I and II the conditions are the same. It can be seen that the *EC*_50_ and *SDM* values between diffusion and reference assays do not considerably differ (Table S4) and thus no substantial improvement regarding sensitivity could be achieved. The concentration-response curves of hsa-miR-30a-3p and miR-93-5p for the standard reference assay are shown in Fig. S7 and for the adapted reference assay, the diffusion assay I and II in Fig. S8. Both miRNAs show comparable results to miR-21-5p.

